# Cryptococcosis, tuberculosis, and a kidney cancer fail to fit the atherosclerosis paradigm for foam cell lipid content

**DOI:** 10.1101/2023.06.08.542766

**Authors:** Valentina Guerrini, Brendan Prideaux, Rehan Khan, Selvakumar Subbian, Yina Wang, Evita Sadimin, Siddhi Pawar, Rahul Ukey, Eric A. Singer, Chaoyang Xue, Maria Laura Gennaro

**Author notes:** Boehringer Ingelheim Animal Health, Ames, IA 50010 (VG); City of Hope National Medical Center, Duarte, CA 91010 (ES); Division of Urologic Oncology, The Ohio State University Comprehensive Cancer Center, Columbus, OH 43221 (EAS). First authorship (Valentina Guerrini, Brendan Prideaux, Rehan Khan).

## Abstract

Foam cells are dysfunctional, lipid-laden macrophages associated with chronic inflammation of diverse origin. The long-standing paradigm that foam cells are cholesterol-laden derives from atherosclerosis research. We previously showed that, in tuberculosis, foam cells surprisingly accumulate triglycerides. Here, we utilized bacterial (*Mycobacterium tuberculosis*), fungal (*Cryptococcus neoformans*), and human papillary renal cell carcinoma (pRCC) models to address the need for a new explanation of foam cell biogenesis. We applied mass spectrometry-based imaging to assess the spatial distribution of storage lipids relative to foam-cell-rich areas in lesional tissues, and we characterized lipid-laden macrophages generated under corresponding *in vitro* conditions. The *in vivo* data and the *in vitro* findings showed that cryptococcus-infected macrophages accumulate triglycerides, while macrophages exposed to pRCC- conditioned-medium accumulated both triglycerides and cholesterol. Moreover, cryptococcus- and mycobacterium-infected macrophages accumulated triglycerides in different ways. Collectively, the data show that the molecular events underlying foam cell formation are specific to disease and microenvironment. Since foam cells are potential therapeutic targets, recognizing that their formation is disease-specific opens new biomedical research directions.

---

Chronic inflammation of infectious and non-infectious origin is often associated with the presence of foam cells, lipid-laden macrophages that exhibit impaired immune function and can contribute to pathogenesis ^1^. Foam cells (or foamy macrophages) form when, due to dysregulated metabolism, lipids accumulate beyond the homeostatic capacity of macrophages. The lipids are stored as droplets that confer a foamy appearance to the macrophages ^2^. Our understanding of foam cell biology is based on studies of atherogenesis, a disease in which uptake of normal and proinflammatory lipoproteins by macrophages in the arterial wall leads to imbalanced cholesterol metabolism and formation of cholesterol-laden foam cells ^3^. Accumulation of foam cells in the arterial intima leads to chronic inflammation, cell death, and tissue necrosis ^3^. A similar situation is observed in tuberculosis, a chronic inflammatory disease of the lung caused by *Mycobacterium tuberculosis*. In the tuberculous lung lesions, which are called granulomas, the presence of tissue necrosis is associated with foam cell accumulation^1^. Indeed, foam cells are a hallmark of both the atherosclerotic plaque and the necrotizing tuberculous granuloma ^3,4^. Thus, we were surprised to find that tuberculous foam cells are enriched in triglycerides rather than cholesterol ^5^, as was the expectation derived from atherogenesis. Whether tuberculosis is an outlier or whether it represents a common situation requiring abandonment of the atherogenesis paradigm is unknown.

To test the hypothesis that foam cell biogenesis is disease-specific, we began a study of foam cells associated with the fungal infection cryptococcosis and with papillary renal cell carcinoma, a cancerous condition. Cryptococcosis is a clinically heterogeneous disease caused by the fungal pathogen *Cryptococcus neoformans*. It affects the lung and other organ systems, including the central nervous system, particularly in immunocompromised individuals ^6^. Foamy macrophages are observed in human tissue biopsies from pulmonary and extrapulmonary cryptococcosis ^7–9^ and in the lungs of infected mice ^10^. Papillary renal cell carcinoma (pRCC), as well as several forms of cancer of many organ systems (liver, lung, colon/rectum, and kidney), also have associated foamy macrophages ^11–16^. The nature of storage lipids is unknown for these pathologies.

In the present work, we assessed the spatial distribution of foamy macrophages and storage lipids in *C. neoformans*-infected murine lungs and in human pRCC specimens. We then analyzed lipid content and the transcriptional program of lipid-laden macrophages generated under *in vitro* conditions that corresponded to these two diseases. The data establish that foam cell formation varies with disease context. We can no longer base our understanding of foam cell biogenesis only on atherogenesis studies. Expanding our view of foam cell biogenesis is expected to provide new targets for therapeutic intervention into diseases -- such as atherosclerosis, tuberculosis, multiple sclerosis, and certain cancers -- in which foam cell appearance is associated with poor clinical outcome.

## Materials and Methods

The supplementary materials include the description of the materials and methods utilized to generate, culture, infect and/or treat primary human monocyte-derived macrophages; to perform measurements of neutral lipid content; RNA extraction and bulk RNA sequencing with the associated statistical analyses; to conduct mouse infections with *C. neoformans*; to obtain and process cryptococcus-infected murine lung specimens and human cancerous kidney surgical resections; to conduct histopathological analysis of the lesional tissues; to perform matrix-assisted laser desorption/ionization mass spectrometry for the analysis of the spatial distribution of triglycerides and cholesteryl esters in lesional tissues.

## Results

### Foamy macrophages cluster peri- or extra-lesionally and associate with triglycerides in *C. neoformans*-infected murine lungs

Foamy macrophages form during pulmonary and extrapulmonary cryptococcal infection ^7,8^. We used a model of pulmonary cryptococcosis with C57BL/6 mice to assess the spatial relationship between foam cells and neutral lipids [triglycerides (TAG) and cholesteryl esters (CE)] in infected lungs. At 7 days post intranasal infection, infected mouse lungs exhibited several granulomatous nodular lesions visible at low magnification (**Fig. S1**). The lesions consisted of large aggregates of fungal cells surrounded by inflammatory infiltrates comprised mostly of polymorphonuclear cells, macrophage and lymphocyte aggregates, and epithelioid cells (**Fig. 1A**). Macrophages contained small nuclei and cytoplasmic lipid droplets giving them a foamy/bubbly appearance. These foam cells tended to form clusters in peri- or extra-lesional areas of the infected lung foci [see hematoxylin and eosin (H&E)-stained lung slices in **Fig. 1BC**]. When we used matrix-assisted laser desorption/ionization mass spectrometry (MALDI) imaging of sections adjacent to those used for H&E staining, we detected multiple TAG and CE species in the infected lungs (**Table S1**). All CE species localized in the fungus-rich lesions (e.g., compare H&E staining and MALDI imaging for CE 16:0 in **Fig. 1C**). In contrast, TAG species were distributed throughout the lung tissue, with some species, such as TAG 46:0, being more prominently found within the lesions and others, such as TAG 50:1, found extra-lesionally (**Fig. 1C**, with corresponding ion counts in **Fig. 1D**) (see **Fig. S2** for uninfected control tissue). Localization of some TAG species and CE species in the fungus-rich lesions is consistent with the presence of both TAG and sterols in fungal cells ^17,18^. In addition, the spatial distribution of some TAG species, such as TAG 50:1 in **Fig. 1C**, which was present throughout the tissue but approximately two-fold lower in the fungus-rich lesions, was characteristic of these foam cells (compare H&E staining and MALDI imaging in **Fig. 1C**). Thus, the pathogen cells appear to contain both TAG and CE, while the *Cryptococcus*-induced foam cells are TAG, not CE, enriched.

**Fig. 1.**
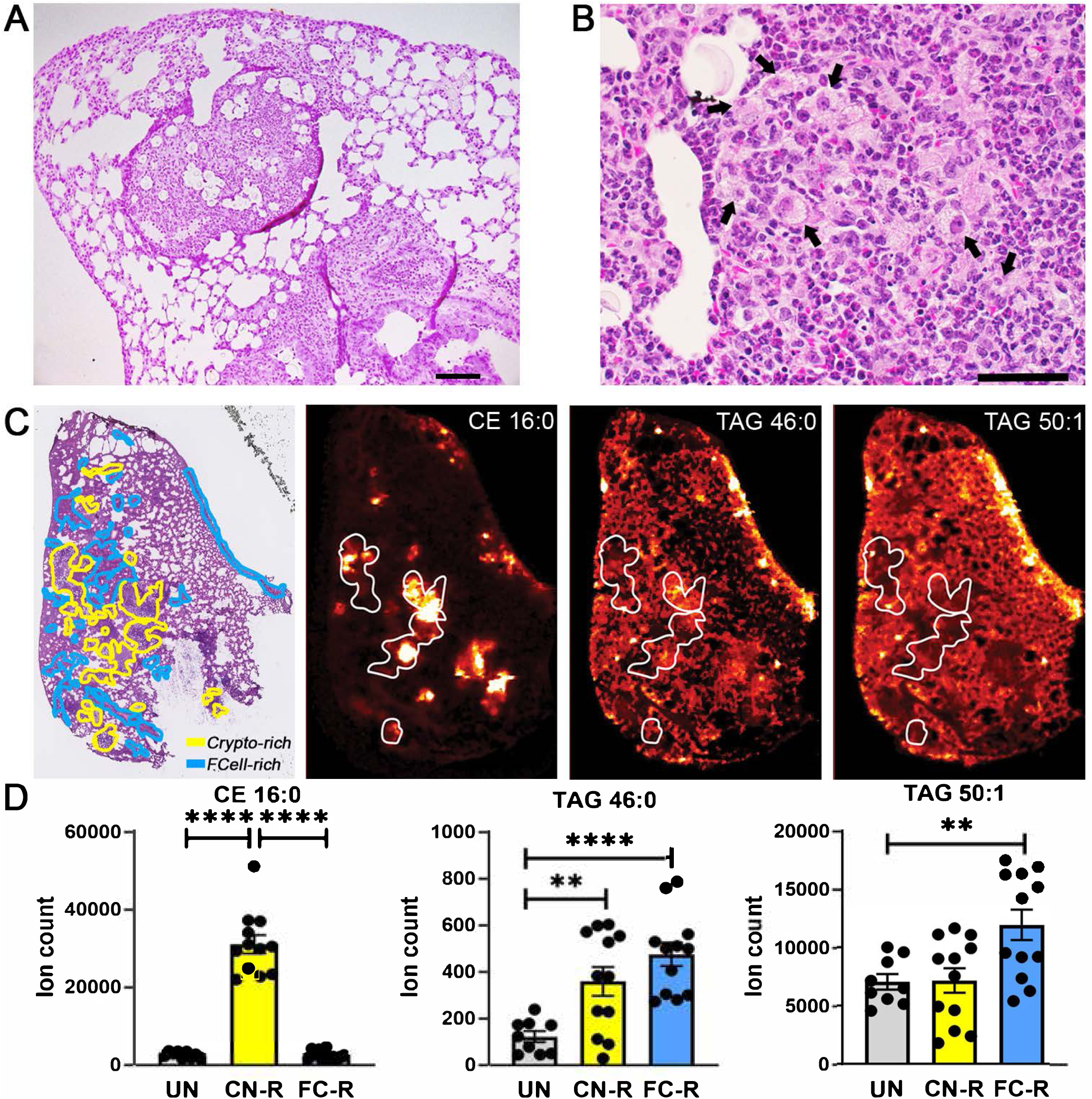
Spatial distribution of foam cells and storage lipids in *C. neoformans*- infected murine lungs. **A-B.** <u>H&E staining of formalin-fixed, paraffin-embedded lung</u> <u>sections from *C.*</u> *<u>neoformans</u>* <u>H99-infected mice</u>. Images were photographed at (A) 100x magnification; scale bar is 100 μm; and (B) 400x magnification; scale bar is 10 μm. Black arrows indicate foam cells. **C.** <u>MALDI imaging of representative CE and TAG</u> <u>species in infected lung sections</u>. The left panel shows H&E staining of infected tissue sections (scale bar is 2 mm). The yellow lines delineate areas enriched in fungal cells (Crypto-rich), while the blue lines define areas enriched in foam cells (FCell-rich). The three additional panels show MALDI imaging of storage lipids in lung sections contiguous to those used for H&E staining. Representative species are shown: CE (16:0) [M+K]^+^ *m/z* 663.48 and TAG (46:0) [M+K]^+^ *m/z* 817.669 signals tend to correspond to cryptococci-enriched areas, while TAG (50:1) [M+K]^+^ *m/z* 871.716 tends to be reduced in those same areas. Areas delimited by white lines correspond to some cryptococci-enriched areas in the H&E-stained section. Corresponding images of uninfected lung sections are shown in *Fig. S2*. **D.** <u>Quantification of CE and TAG MALDI</u> <u>imaging intensity (expressed as ion count)</u>. Quantification of lipid species was performed in uninfected tissue and in fungus-rich (CN-R) and foam-cell-rich (FC-R) areas of the infected tissue. Mean and SEM of 9 sections from three uninfected animals and 12 sections from four infected animals (3 sections per animal) are shown. **, *p* < 0.05; ****, *p* < 0.001 (unpaired *t*-test).

### In papillary renal cell carcinoma foamy macrophages preferentially associate with CE- and TAG-enriched kidney areas

Papillary renal cell carcinoma (pRCC) is useful for studies of foam cell biogenesis in a cancer context, because foam cells are a frequent histopathologic finding ^15,16^. To characterize pRCC-associated foam cells, we used specimens obtained from patients who underwent partial or radical nephrectomy and performed MALDI imaging and H&E staining on adjacent sections of the resected tissues. The foamy macrophages, which are characterized morphologically by foamy/bubbly cytoplasm and small nuclei, were interspersed throughout the inter-tumoral stroma. Nine CE species were detected in the pRCC tissues. Most were distributed throughout the tissue, but their localization varied with the degree of saturation of the esterified fatty acid (**Fig. S3**, **Table S1**). In particular, the two monounsaturated species (CE 16:1 and CE 18:1), which were the most abundant in the tissues, were present as highly localized, intense signals (CE 16:1 tissue localization is shown in **Fig. 2A**). In contrast, TAG species yielded localized signals that were similar for all detected TAGs (**Fig. 2B** shows the distribution of TAG 52:2, which is representative of all TAG species; see **Fig. S4** for other TAG species). H&E staining revealed that the intense, localized CE signals correspond to large foam cell aggregates (**Fig. 2CEG** show one such area at increasing magnification). In contrast, the TAG signals corresponded to tissue regions containing large numbers of foam cells interspersed among cancer cells (**Fig. 2DFH** show a representative area at increasing magnification). Since CE species were detected throughout the tissue (**Fig. S3**), the TAG-rich regions also contained CE, albeit at lower levels than in the large foam cell aggregates shown in the left panels of **Fig. 2**. Bioptic tissues collected from two additional patients showed similar differential distribution of TAG and CE species in pRCC foam cells, with more intense TAG signals associated with large necrotic regions (**Fig. S5**). In summary, MALDI imaging showed associations between foam-rich areas with TAG species, CE species, or both. Thus, pRCC illustrates a class of foam cells in which both CE and TAG are present.

**Fig. 2.**
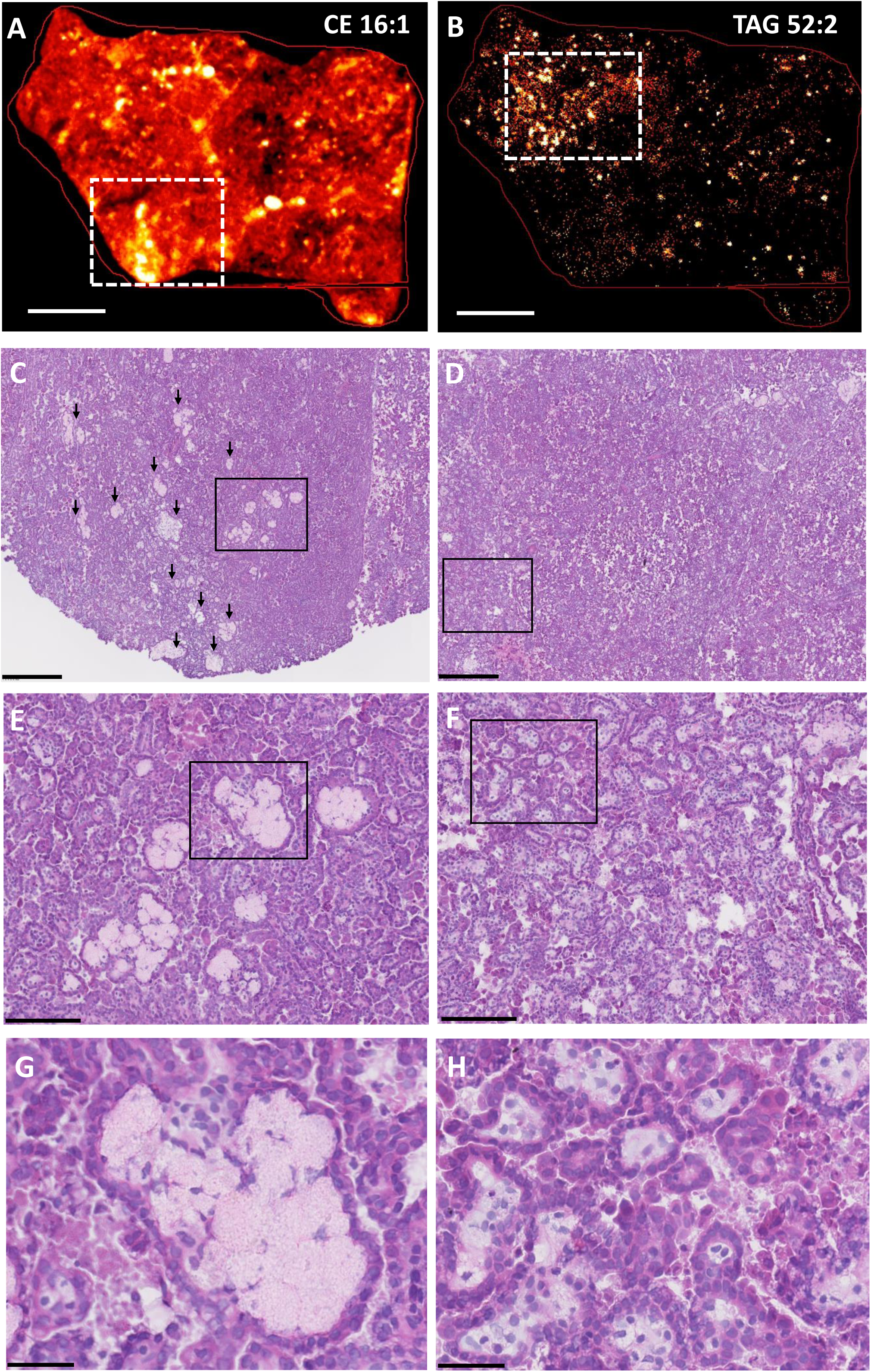
Spatial distribution of foam cells and storage lipids in papillary renal cell carcinoma (pRCC). **A-B.** <u>MALDI imaging of representative CE and TAG species in</u> <u>pRCC resected tissues</u>. MALDI-2 MSI ion distribution for CE 16:1 [M+K]^+^ *m/z* 661.532 (panel A) and TAG 52:2 [M+K]^+^ *m/z* 897.731 (panel B) are shown in frozen pRCC tissue sections; scale bar is 3 mm. The white rectangles delineate areas of high signal intensity that are magnified in the corresponding histology panels C-D. **C-H.** <u>H&E</u> <u>staining of frozen pRCC tissue sections</u>. Serial sections to those used for MALDI imaging were H&E stained. Each column corresponds to the top MALDI image (left panels, CE 16:1; right panels, TAG 52:2). The black box in each row marks the area of tissue shown at higher magnification in the corresponding panel below. **C-D**. scale bar is 500 mm. Black arrows in panel C mark large foam cell aggregates. The black box in C marks an area enriched for foam cell aggregates, which are further magnified in panel E. The black box in D marks an area enriched for foam cells interspersed among tumor cells, which is further magnified in panel F. **E-F.** scale bar is 150 mm. The black boxes in these panels mark areas further magnified in panels G and H, respectively. **G-H.** scale bar is 50 mm. Panel G shows a foam cell aggregate; panel H shows foam cells interspersed among tumor cells.

### *Cryptococcus neoformans* infection induces accumulation of triglyceride-rich lipid droplets in macrophages via an mTORC1-independent pathway

MALDI imaging provides information about the spatial distribution of analytes in tissues, but it does not have the single-cell resolution needed to precisely assign a particular neutral lipid to a specific cell type. Thus, we utilized an in vitro infection model to study neutral lipid accumulation in macrophages infected with *C. neoformans*. When we infected primary human monocyte-derived macrophages (MDM) with mCherry-expressing *C. neoformans* H99 and quantified lipid droplet content by imaging flow cytometry, we observed significant lipid droplet accumulation in infected macrophages (3.5-fold increase relative to uninfected cells) (**Fig. 3AB**). Lipid-droplet-enriched macrophages in the infected culture wells included both those containing fungal cells and those that did not (**Fig. 3A** and quantitative data in **Fig. S6**), indicating that lipid droplet formation does not require internalization of fungal cells.

**Fig. 3.**
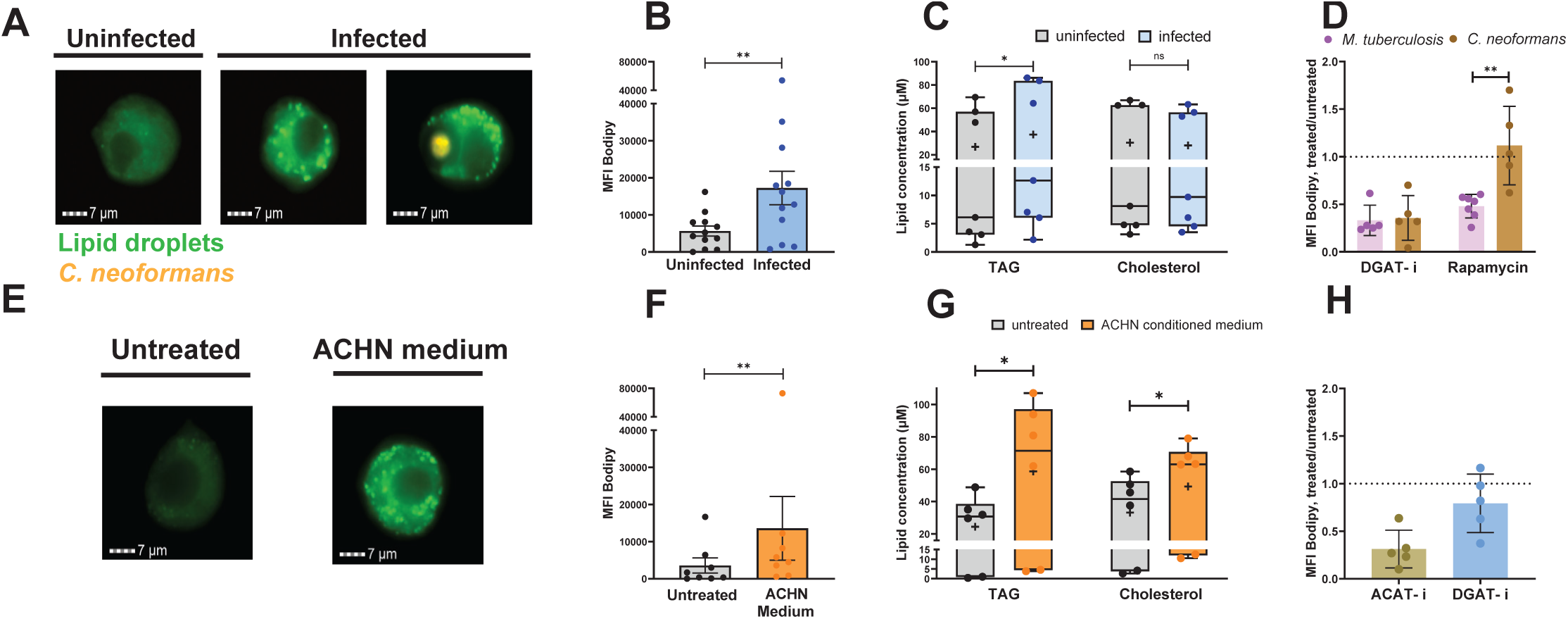
Characterization of lipid droplets induced in human primary macrophages by *C. neoformans* infection and by exposure to conditioned medium from ACHN cell cultures, and comparisons between *C. neoformans*- and *M. tuberculosis*- induced effects. <u>Panels A-D</u> show data obtained with monocyte-derived macrophages (MDM) infected with *C. neoformans* for 24 h (MOI = 4). Panel D also includes cells infected with *M. tuberculosis* for 24 h (MOI = 4). In all bar graphs, each dot corresponds to one human donor. **A.** <u>Lipid droplet imaging</u>. Representative images of MDM uninfected (leftmost panel), infected with mCherry-tagged *C. neoformans,* and stained with Bodipy 493/503 (neutral lipid dye, green fluorescence) were acquired by imaging flow cytometry at 24 h post-infection. The two rightmost panels show macrophages in the infected culture wells carrying and not carrying intracellular fungi (orange fluorescence). **B.** <u>Lipid droplet content</u> was expressed as median fluorescence intensity (MFI; +/- SD) of Bodipy 493/503. **C.** <u>Neutral lipid measurements</u>. TAG and cholesterol were measured in uninfected and infected cells, as indicated, using a commercially available kit. The box plots show lower quartile, median, and upper quartile of the distribution of multiple donors. The whiskers represent minimum and maximum values. The plus symbol indicates the mean. ns, non-significant; *, *p* < 0.05 (paired *t*-test). **D.** <u>Effect on lipid droplet content of treatment with chemical inhibitors</u>. DMSO (vehicle control), 0.4 nM rapamycin (mTORC1 inhibitor), or 30 nM DGAT-1 inhibitor (DGAT-i) (A922500; PubChem CID: 24768261) were added for the duration of infection. Lipid droplet content was quantified by imaging flow cytometry and expressed as Bodipy MFI, as in panel A. Results are shown as ratios of Bodipy MFI of drug-treated to vehicle- treated infected cells. Mean and SD are shown. ns, non-significant; **, *p* < 0.01 (unpaired *t*-test). <u>Panels E-H</u> show data obtained with MDM left untreated and treated with ACHN-conditioned medium for 7 days. **E.** <u>Lipid droplet imaging</u>. Cells were stained with Bodipy 493/503 at the end of treatment and images were acquired by imaging flow cytometry, as in panel A. **F.** <u>Lipid droplet content</u> was expressed as MFI of Bodipy 493/503, as in panel B. **G.** <u>Neutral lipid measurements</u>. TAG and cholesterol were measured in untreated and ACHN-medium-treated cells and data expressed as described in panel C. **H.** <u>Effect on lipid droplet content of treatment with chemical</u> <u>inhibitors</u>. ACHN-medium-treated MDM were treated with DMSO (vehicle control), 90 n DGAT-1 inhibitor (DGAT-i) (A922500; PubChem CID:24768261), or 10 μM ACAT inhibitor (ACAT-i) (CAS 615264-52-3; PubChem CID:10019206) for 7 days. Results are shown as ratios of Bodipy MFI of drug-treated to vehicle-treated cells, as in Panel D.

When we measured storage lipid content in *C. neoformans*-infected cells by an enzymatic assay, we found that infection increased the content of intracellular TAG but not cholesterol derivatives (**Fig. 3C**). Moreover, lipid droplet accumulation in *C. neoformans*-infected cells was essentially abrogated by treatment with A922500, an inhibitor of diglyceride acetyl transferase (DGAT), the enzyme that catalyzes the conversion of diglycerides to triglycerides (**Fig. 3D**). This finding supports the conclusion that *C. neoformans*-induced lipid droplets are TAG enriched, as also seen with *M. tuberculosis* infection (^5^ and **Fig. 3D**). This result agrees with the above observation that *C. neoformans* lung infection is associated with TAG-enriched foam cells.

Our previous work showed that the accumulation of TAG-rich lipid droplets in macrophages infected with *M. tuberculosis* requires signaling by mechanistic target of rapamycin complex 1 (mTORC1), as it is inhibited by rapamycin treatment ^5^. Unlike the *M. tuberculosis* case, however, rapamycin had no effect on lipid droplet accumulation in *C. neoformans-*infected macrophages (**Fig. 3D**). Thus, even though *M. tuberculosis* and *C. neoformans* both induce accumulation of TAG-rich lipid droplets, the two pathogens do so by utilizing different signaling pathways. These data emphasize the diversity of lipid droplet formation.

### Factor(s) released by a papillary renal cell carcinoma-like cell line induce macrophage accumulation of both triglycerides and cholesteryl esters

We next investigated the effects of pRCC on storage lipid accumulation in macrophages in vitro by exposing human macrophages to cell-free conditioned medium from cultures of the ACHN cell line, which is derived from a human renal cell carcinoma and exhibits pRCC features ^19^. ACHN-medium-treated macrophages also exhibited lipid droplet accumulation (**Fig. 3EF**), in agreement with previous observations ^13^. Storage lipid analysis by enzymatic assays showed that lipid droplet accumulation in ACHN-medium-treated macrophages correlated with increased levels of both TAG and cholesterol (**Fig. 3G**). Moreover, the lipid droplet content of these macrophages decreased upon treatment with CAS 615264-52-3, a chemical inhibitor of acyl-coenzyme A:cholesterol acyltransferase (ACAT), the enzyme that converts cholesterol to cholesteryl esters, and, to some extent, also with the DGAT inhibitor A922500 (**Fig. 3H**). Together, these results are consistent with the conclusion of the above analysis of kidney bioptic tissues that pRCC-associated foam cells contain both TAG and CE, suggesting yet another context-specific mechanism of foam cell formation.

### Transcriptomics identify condition-specific molecular events underlying macrophage lipid droplet accumulation

We utilized transcriptomics to investigate the pathways underlying neutral lipid accumulation in macrophages from the same donors that were infected with *M. tuberculosis* and *C. neoformans,* and treated with ACHN conditioned medium. When we analyzed the Gene Ontology (GO) annotations related to metabolic processes, we found that the most informative signatures of macrophage metabolic reprogramming associated with TAG accumulation were derived from the top-ranked downregulated pathways in *M. tuberculosis*-infected macrophages, which included lipid catabolism, fatty acid oxidation, oxidative phosphorylation, and electron transport chain (**Fig. 4A**), and from the top-ranked upregulated pathways in *C. neoformans*-infected macrophages and ACHN-medium-treated macrophages, which were both enriched for glycolysis (**Fig. 4BC**) (the remaining pathway analysis results are found in **Fig. S7-S8**).

**Fig. 4.**
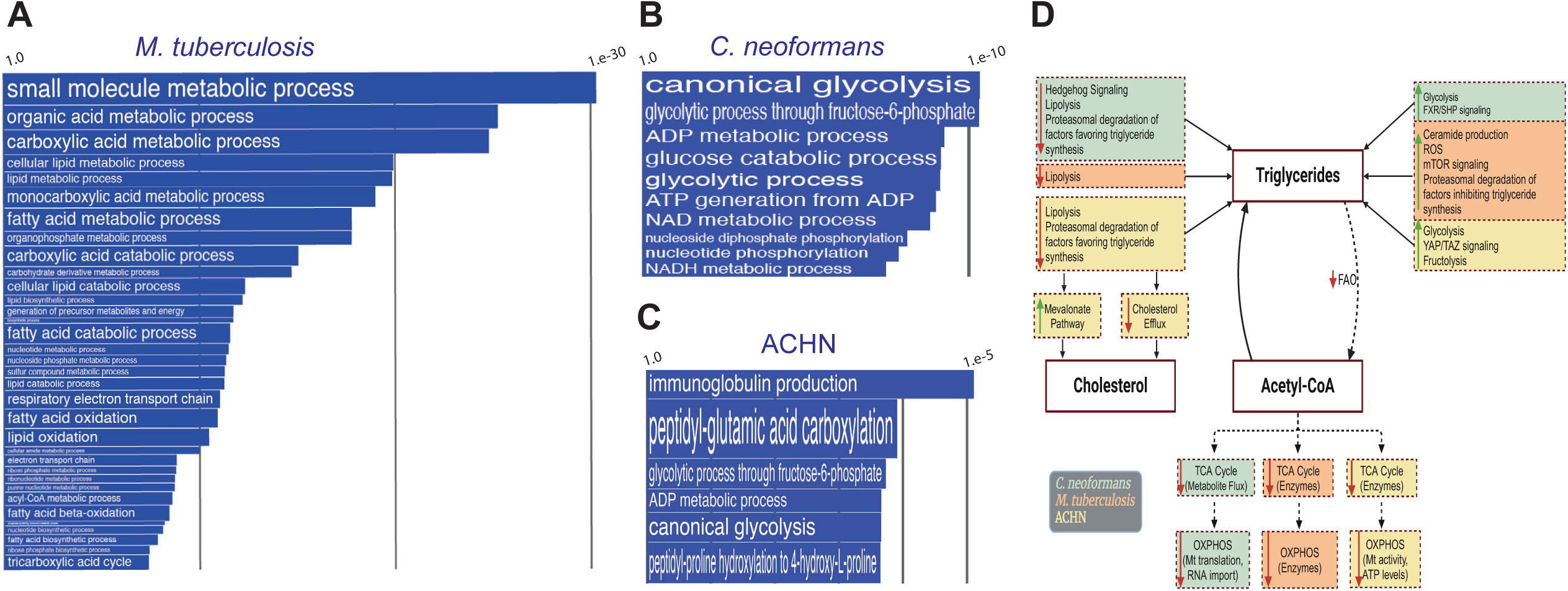
Transcriptomics analysis of monocyte-derived macrophages infected with *M. tuberculosis* and *C. neoformans*, and exposed to ACHN conditioned medium. Cells from the same donors were used across experimental conditions, as indicated in Fig. 3 legend. RNA was isolated and subjected to RNA-seq analysis, as described in *Methods*. **A-C.** <u>Pathway analysis</u>.The panels show Gene Ontology (GO) annotations related to metabolic processes that were (**A**) downregulated in *M. tuberculosis*-infected MDM, (**B**) upregulated in *C. neoformans*-infected MDM, and (**C**) upregulated in ACHN- medium-treated MDM, relative to control cells. The differential expression between sample classes (infected vs uninfected or treated vs untreated) was tested with coincident extreme ranks in numerical observations (CERNO). Pathways were selected using a cutoff false discovery rate of 0.05; the *p*-values for these pathways are plotted onto the x-axis. To represent effect size, pathway gene sets containing fewer genes were given greater bar height/font size than were larger sets that yielded similar *p* values. For visualization purposes, only top ranking annotations are shown. In panel C, it is noted that immunoglobulin production by macrophages in the context of tumor microenvironments has been described ^37^ - it is not discussed as it does not appear relevant to the topic of our report. Additional pathway analysis data are shown in *Fig. S7* and *Fig. S8*. **D.** <u>Comparative summary of differentially regulated pathways associated</u> <u>with neutral lipid accumulation in the experimental three conditions tested</u>. The figure shows altered key cellular functions and signaling pathways, as revealed by transcriptomics, that may contribute to accumulation of triglycerides in the three experimental conditions studied in this report, and to cholesterol accumulation in ACHN- medium treated cells. Differentially expressed pathways in each experimental condition (relative to controls) are shown in boxes that are color-coded with respect to the experimental condition (*C. neoformans*, light green; *M. tuberculosis*, light pink; ACHN, light yellow). Green arrows, upregulation; red arrows, downregulation. The straight black arrows toward “triglycerides” and “cholesterol” signify lipogenic processes. The arrows originating from or directed to “Acetyl-CoA” signify increased (solid line) or decreased (broken lines) relative availability of acetyl-CoA. Additional relationships between oxidative phosphorylation/mitochondrial function and triglyceride accumulation are not indicated. FAO, fatty acid oxidation; OXPHOS, oxidative phosphorylation; Mt, mitochondrial.

Additional insight was derived from analyses at the gene level. In *M. tuberculosis*- infected macrophages, the top three downregulated metabolic genes encoded: (i) acyl-CoA synthase (ACSM5), (ii) carnitine octanoyl transferase (CROT), which converts acyl-CoA to acyl-carnitine, a step required for transport across the mitochondrial membrane, and iii) aldehyde hydrogenase (ALDH3A2), which oxidizes fatty aldehydes to fatty acids (**Table S2** and *Supplementary text*). Downregulation of these genes likely leads to defective fatty acid oxidation. The top-ranking upregulated pathways and individual genes in *M. tuberculosis*-infected macrophages related to ubiquitination processes, which presumably result in substantial macrophage proteome remodeling in response to infection, including various lipid metabolism regulators (**Table S2** and *Supplementary text*).

In *C. neoformans* infection, the top five upregulated metabolic genes all encoded glycolytic enzymes, i.e., hexokinase 2 (HK2), fructose-bisphosphate aldolase C (ALDOC), glyceraldehyde 3-phosphate dehydrogenase (GAPDH), phosphoglycerate kinase 1 (PGK1), and phosphopyruvate hydratase (ENO2) (**Table S3**). Among the top-ranking downregulated genes in *C. neoformans*-infected macrophages featured indicators of reduced mitochondrial functions, including downregulation of polyribonucleotide nucleotidyl transferase 1 (PNPT1) and a glutaminyl-tRNA amidotransferase subunit 1 (QRSL1) (**Table S3**). The PNTP1 product regulates mitochondrial homeostasis and the abundance of electron transport chain components^20^. Missense mutations in the human QRSL1 locus have been associated with defects in oxidative phosphorylation ^21^. Moreover, the downregulation of anaphase promoting complex subunit 7 (**ANAPC7**) may also lead to triglyceride accumulation through activation of farnesoid X receptor (FXR) signaling ^22^ (**Table S3** and *Supplementary text*). Together, the transcriptomics data strongly suggest that the accumulation of TAG results from increased glycolysis in *C. neoformans*-infected macrophages and decreased lipid catabolism, TCA cycle, and oxidative phosphorylation by different molecular modalities in *M. tuberculosis*- and *C. neoformans*-infected macrophages (**Fig. 4D**).

In ACHN-medium-treated macrophages, the top-ranking upregulated metabolic genes encoded glycolytic enzymes, i.e., hexokinase 3 (HK3), phosphoglycerate kinase (PGK1), and a phosphopyruvate hydratase (ENO2) (**Table S4**). In addition, triosephosphate isomerase (TPI1) contributes to the conversion of dihydroxyacetone phosphate to glyceraldehyde 3-phosphate, favoring triglyceride synthesis (**Table S4,** and *Supplementary text*). Thus, similarities exist with the top-ranking upregulated metabolic functions of *C. neoformans*-infected cells. We also found gene markers of reduced TCA cycle in ACHN-medium-treated macrophages, including upregulation of adenylate kinase 4 (AK4), a key metabolic regulator that increases glycolysis and inhibits the TCA cycle and oxidative phosphorylation ^23^, and downregulation of PPARGC1A (**Table S4**). The latter gene encodes PGC-1α, a master regulator of energy metabolism that promotes fatty acid oxidation and the TCA cycle, thereby decreasing TAG storage ^24^ (**Table S4** and *Supplementary text*). Together, increased glycolysis and reduced TCA cycle would result in routing pyruvate towards de novo lipogenesis and, consequently, explain lipid droplet accumulation in macrophages. Additional top-ranking downregulated genes that contribute to lipid accumulation in ACHN-medium-treated cells are listed in **Table S4** and their function discussed in the *Supplementary text*. Overall, exposure to ACHN conditioned medium is associated with yet other, distinctive modalities of increased glycolysis, decreased lipid catabolism and degradation, and decreased TCA cycle and oxidative phosphorylation, leading to triglyceride accumulation (**Fig. 4D**).

Additional gene-level expression analyses of the three experimental conditions examined further revealed molecular events that may lead to TAG accumulation. In *M. tuberculosis* infection, these include downregulation of lipolytic genes, upregulation of sirtuins and sirtuin-stabilizing functions, and expression changes in genes signifying increased production of ceramide and altered cellular redox (see **Table S2** and *Supplementary text*). In *C. neoformans* infection, additional indicators of metabolic remodeling toward TAG biosynthesis included (i) upregulation of genes for the production of dihydroxyacetone phosphate, which can be routed toward TAG biosynthesis, (ii) upregulation of hexokinase (HK2) and lactate dehydrogenase (LDHA), which indirectly inhibit lipolysis, and (iii) downregulation of AMP-activated protein kinase (AMPK), which inhibits de novo biosynthesis of fatty acids and stimulates fatty acid oxidation ^25^ (see **Table S3** and *Supplementary text*). In ACHN-medium treated macrophages, markers of TAG accumulation included the increased expression of genes associated with or regulated by YAP/TAZ signaling, which regulates cancer cell metastasis and metabolic reprogramming, including lipid metabolism ^26^. These genes include TEAD transcription factors, the perilipin PLIN5, and the fructose transporter SLC2A5 (increased fructose uptake may lead to lipogenesis via fructolysis) (**Table S4** and *Supplementary text*). Increased YAP/TAZ signaling is also supported by upregulation of AK4 and downregulation of phospholipase D family member 6 (PLD6); both gene expression changes might result in decreased activity of AMPK, which inhibits YAP/TAZ ^27^ (**Table S4** and *Supplementary text*). These additional observations further point to different molecular processes underlying triglyceride accumulation in macrophages infected with *M. tuberculosis* or *C. neoformans*, or treated with ACHN conditioned medium (**Fig. 4D**).

### Gene expression markers of mTORC1 signaling in *M. tuberculosis*-infected macrophages

The transcriptomics data also shed light on the requirement in *M. tuberculosis* infection for signaling by mTORC1 (**Fig. 2D**), which is lipogenic in multiple ways ^28^. *M. tuberculosis*-infected macrophages downregulated the TP53 gene and upregulated TP53-specific E3 ligases that target this factor for proteasomal degradation (see **Table S2** and *Supplementary text*). Decreased activity of TP53 correlates well with increased mTORC1 signaling, since TP53 induces expression of Deptor (**Table S2**) and leads to activation of AMPK, two factors that inhibit mTORC1 ^29,30^. Thus, the gene expression profiles are in agreement with the rapamycin sensitivity of lipid droplet accumulation in *M. tuberculosis*-infected macrophages.

### Gene expression markers of cholesterol dysregulation in macrophages exposed to ACHN conditioned medium

The ACHN-medium-treated macrophages also exhibited gene expression changes associated with dysregulation of cholesterol metabolism. For example, the scavenger receptor CD36, which is a key regulator of cholesterol homeostasis, was downregulated, presumably as a consequence of PPARGC1A downregulation ^24^ (**Table S4**). CD36 induces cholesterol depletion by promoting macrophage cholesterol efflux and proteasomal degradation of HMG-CoA reductase, the rate-limiting enzyme in sterol synthesis ^31^. An additional marker of dysregulated cholesterol homeostasis is the downregulation of adenylate cyclase (ADCY1), which generates cAMP signaling for cholesterol efflux in atherogenic foam cells ^32^. The above-proposed increased YAP/TAZ signaling might also result in cholesterol accumulation, since YAP/TAZ is involved in the metabolism of fatty acids and sterols ^26^. The dysregulation of cholesterol metabolism suggested by gene expression profiling of ACHN-medium-treated macrophages (summarized also in **Fig. 4D**) is consistent with the formation of cholesterol-containing lipid droplets in vitro and with increased cholesterol derivatives in the pRCC-associated foam cells in vivo.

## Discussion

Together with our earlier finding that foam cells in tuberculous lung lesions are triglyceride-enriched ^5^, the in vivo data reported above for pulmonary cryptococcal infection and papillary renal cell carcinoma show that the neutral lipid content of foam cells is disease-context specific. Moreover, the data obtained in vitro with the corresponding experimental models strongly support the notion that the macrophage metabolic reprogramming resulting in lipid droplet accumulation is also condition-specific. That is the case regardless of the chemical nature of the storage lipids accumulated. For example, macrophages infected with *M. tuberculosis* and *C. neoformans* are enriched in TAG, as demonstrated by the drastic lipid droplet decrease caused by pharmacological inhibition of TAG biosynthesis. In both cases, the accumulation of TAG likely results from a switch from oxidative to glycolytic metabolism that includes increased biosynthesis and decreased catabolism of lipids. However, cryptococcosis and tuberculosis differ in the molecular events underlying the metabolic reprogramming of macrophages, as indicated by the different effect of rapamycin on lipid droplet accumulation in the two infections. Still other mechanisms are likely at play in pRCC-associated macrophages, which accumulate neutral lipids by reprogramming both cholesterol and TAG metabolism. Moreover, although gene expression levels do not directly translate into protein levels, protein activity, and metabolic fluxes, the gene expression data presented above clearly imply that the metabolic remodeling leading to neutral lipid accumulation occurs through signaling, regulatory, and effector mechanisms that are specific to each experimental condition. Therefore, the data lead to the conclusion that macrophage foam cells in different diseases vary in storage lipid content and underlying molecular events, even though they may be similar histochemically (lipid droplets consistently confer a “pale bubbly” appearance upon H&E staining) and perhaps even functionally, as discussed below.

A key to understanding foam cell diversity is the biochemical diversity of the microenvironments driving their biogenesis. It is well established that uptake of exogenous lipids can drive foam cell formation. This is the case in atherogenesis, where sequestration of cholesterol-rich lipoproteins in the arterial wall leads to endothelial activation, recruitment of monocytes, and monocyte differentiation into lipoprotein-ingesting phagocytes that become foam cells ^3^. In some cancers, such as colon cancer, the fatty acid-enriched environment induces lipid droplet accumulation in tumor-associated macrophages ^14^. It would be fallacious, however, to associate foam cell formation exclusively with exogenous lipid uptake. For example, in tuberculosis, macrophage lipid accumulation is associated with TLR2 activation by bacterial components ^33,34^ and requires a lipogenic proinflammatory cytokine, TNFα, produced by infected macrophages ^5^. The formation of pRCC-associated foam cells may involve IL-8 and various chemokines produced by cancer cells ^13^. Lung epithelial cells secrete IL-8 in cryptococcosis ^10^, suggesting that this cytokine may favor foam cell formation in this infection. Additional work is needed to identify the exogenous trigger signals (i.e., those generated by microbes, cancer cells, or other cell types) and to determine additional conditions in which foam cells are induced by combinations of exogenous and autocrine/paracrine signals.

It is reasonable to assume that, despite different pathways of biogenesis, the presence of foam cells represents a maladaptive immune response in all pathological contexts they form. Generally, lipid-laden macrophages tend to lose protective immune functions, including phagocytosis, efferocytosis, and autophagy. They can also induce tissue damage, contribute to necrosis, exhibit impaired antimicrobial activity, and even sustain survival of intracellular pathogens (reviewed in ^1^). Indeed, given their contribution to pathogenesis, foam cells have been recognized as targets of pharmacological intervention. Examples are seen with atherosclerosis and some cancers ^14,35^. Moreover, foam cells are often associated with kidney disease, such as focal and segmental glomerulosclerosis and diabetic nephropathy ^36^, in addition to the pRCC investigated in the present work. Their pathophysiological significance in the kidney remains puzzling, and all mechanistic hypotheses on their biogenesis derive from the atherosclerosis literature ^36^. Recognizing that foam cells result not only from macrophage uptake of exogenous lipids but also from stimuli that are microenvironment-specific opens new directions for mechanistic and drug development research of high biomedical significance.

## Acknowledgements

We thank Knowledge Synthesis, Inc. for help with statistical analyses of the transcriptomics data; Karl Drlica for critical comments on the manuscript. This work was funded by NIH grants R01 HL149450, R01 HL149450-S1, R01 AI158911 (M.L.G.), R01AI155647, R01AI169769 (C.X.), P30CA072720 (E.S. and E.A.S.), and a Cancer Prevention and Research Institute of Texas (CPRIT) High Impact High Risk grant RP190669 (B.P.). M.L.G. also acknowledges funding through NJ ACTS (NIH UL1TR003017).

## Abbreviations

ACAT: acyl-coA:cholesterol acyltransferase
CE: cholesteryl esters
DGAT: acyl-CoA:diacylglycerol acyltransferase
H&E: hematoxylin and eosin
MALDI: matrix-assisted laser desorption/ionization mass spectrometry
MDM: monocyte-derived macrophages
pRCC: papillary renal cell carcinoma
TAG: triglycerides.

## Notes

### Competing Interest Statement

The authors have declared no competing interest.

### Summary of Updates

We have changed title; tightened the key message by vastly revising all sections of the manuscript, in particular the results section; added to our MALDI analysis critical papillary renal cell carcinoma surgical resections to increase the number of subjects tested from 1 to 3; and complemented all in vitro work with chemical inhibitors of key enzymes in triglyceride and cholesteryl ester biosynthesis. We also reorganized two of the four main figures, and removed supplementary data that did not contribute significantly to our main message.

