## Supplemental Files for "Cryptococcosis, tuberculosis, and a kidney cancer fail to fit the atherosclerosis paradigm for foam cell lipid content"

**Supplementary Figures**

Fig. S1. Low-magnification images of murine lungs uninfected and infected with *C. neoformans* H99.

Fig. S2. Hematoxylin-eosin staining and MALDI imaging of uninfected murine lung sections.

Fig. S3. Spatial distributions of cholesterol ester (CE) species in papillary renal cell carcinoma (pRCC) (Patient P1).

Fig. S4. Spatial distribution of triglyceride (TAG) species in papillary renal cell carcinoma (pRCC) (Patient P1).

Fig. S5. Spatial distribution of triglyceride (TAG) and cholesteryl ester (CE) species in papillary renal cell carcinoma biopsies collected from patients P2 and P3**.**

Fig. S6. Macrophage lipid droplet accumulation in *C. neoformans­*-infected cells.

Fig. S7. Metabolism-associated pathways (A) upregulated in *M. tuberculosis*-infected human monocyte-derived macrophages and (B) downregulated in *C. neoformans*-infected human monocyte-derived macrophages.

Fig. S8. Metabolism-associated pathways downregulated in ACHN-medium-treated human monocyte-derived macrophages.

**Supplementary Tables**

Table S1. List of triglyceride and cholesteryl ester species detected in tissues by MALDI-MSI.

Table S2a. Top five down- and up-regulated genes in macrophages infected with *M. tuberculosis.*

Table S2b. Genes functionally related to top five down- and up-regulated genes in macrophages infected with *M. tuberculosis.*

Table S3a. Top five down- and up-regulated genes in macrophages infected with *C. neoformans.*

Table S3b. Genes functionally related to top five down- and up-regulated genes in macrophages infected with *C. neoformans.*

Table S4a. Top five down- and up-regulated genes in macrophages treated with ACHN-conditioned medium.

Table S4b. Genes functionally related to top five down- and up-regulated genes in macrophages treated with ACHN-conditioned culture medium.

**Supplementary Text on transcriptomics analyses**

**Materials and Methods**

**Supplementary References**

**Supplementary Figures**

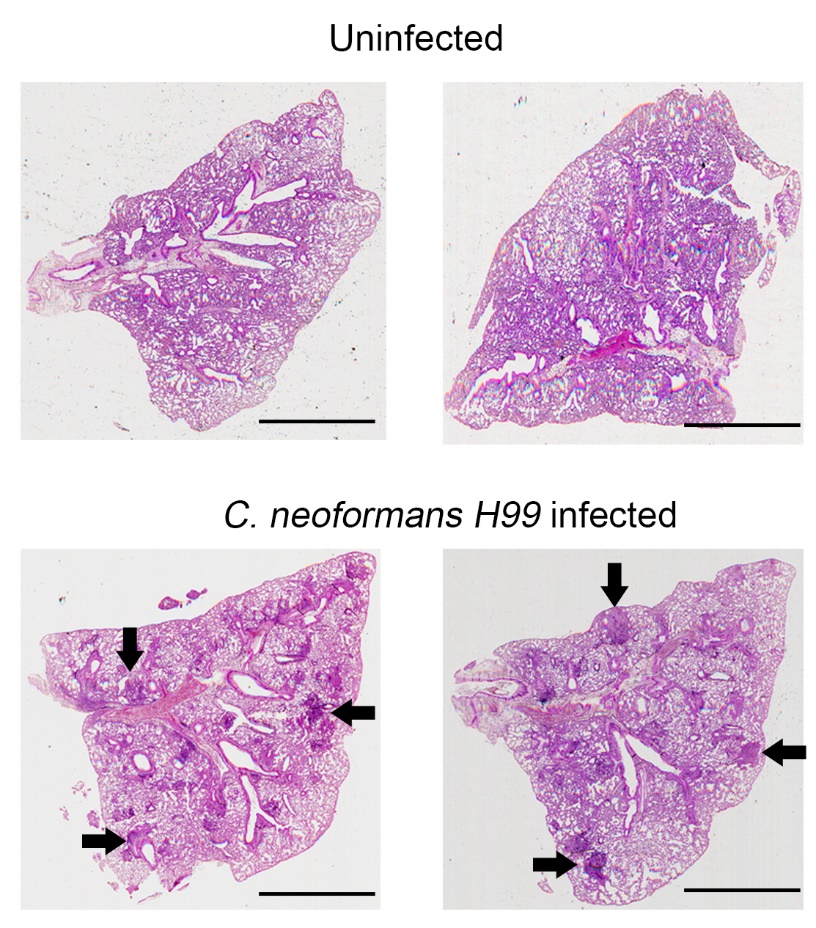

**Supplementary Fig. S1.** **Low-magnification images of murine lungs uninfected and infected with *C. neoformans* H99.** H&E staining of uninfected (top panels) and infected (bottom panels) lung sections. Black arrows show cellular aggregates in the infected tissue. Images were photographed at 20x magnification; scale bar is 2 mm.

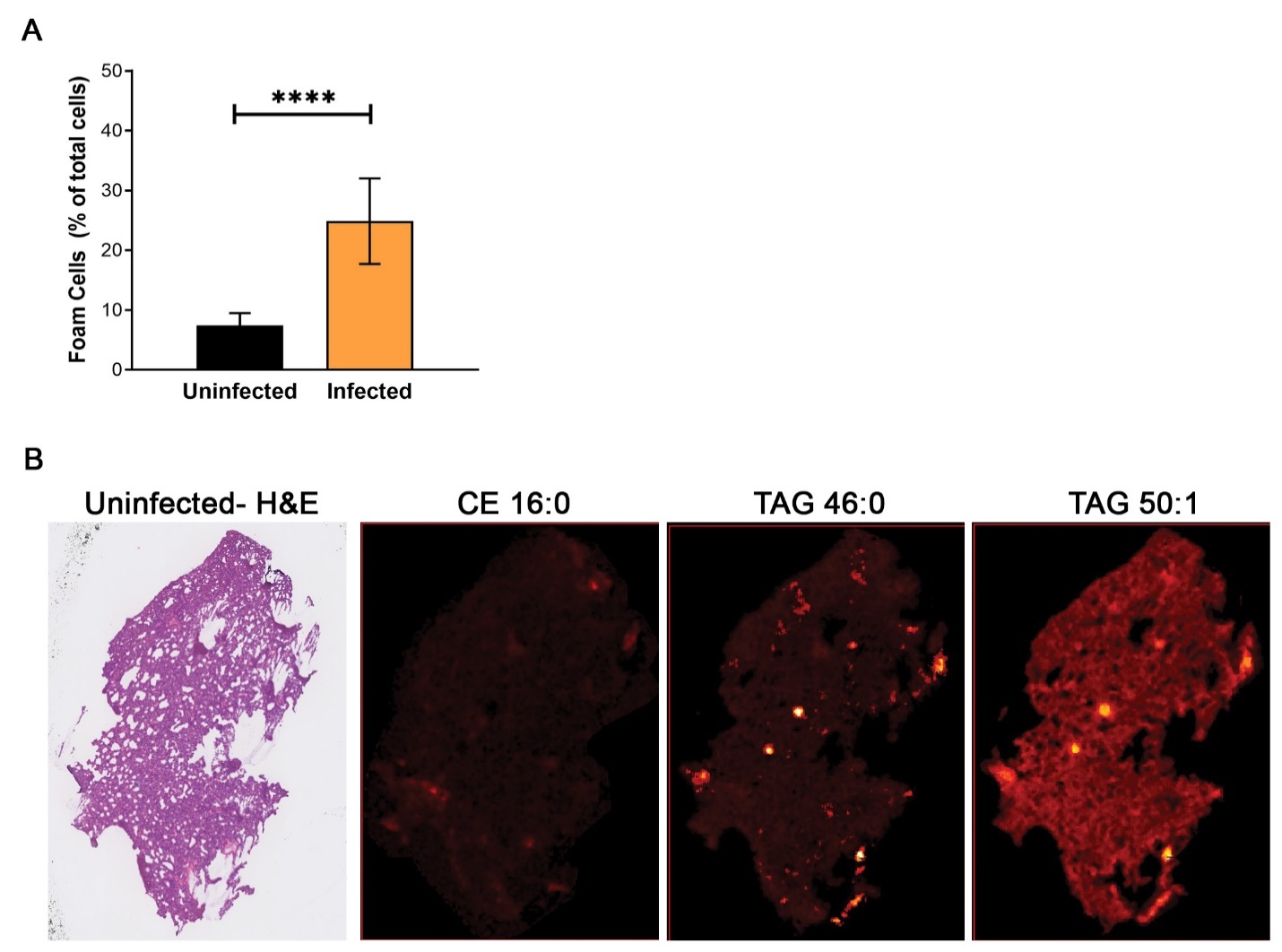

**Supplementary Fig. S2. Hematoxylin-eosin staining and MALDI imaging of uninfected murine lung sections.** The left panel shows H&E staining of uninfected lung sections. The three additional panels show MALDI imaging of storage lipids in lung sections contiguous to those used for H&E staining. Representative species are shown, CE (16:0) and TAG (46:0 and 50:1), corresponding to those shown in Fig. 1C (TAG and CE species detected are listed in **Table S1**).

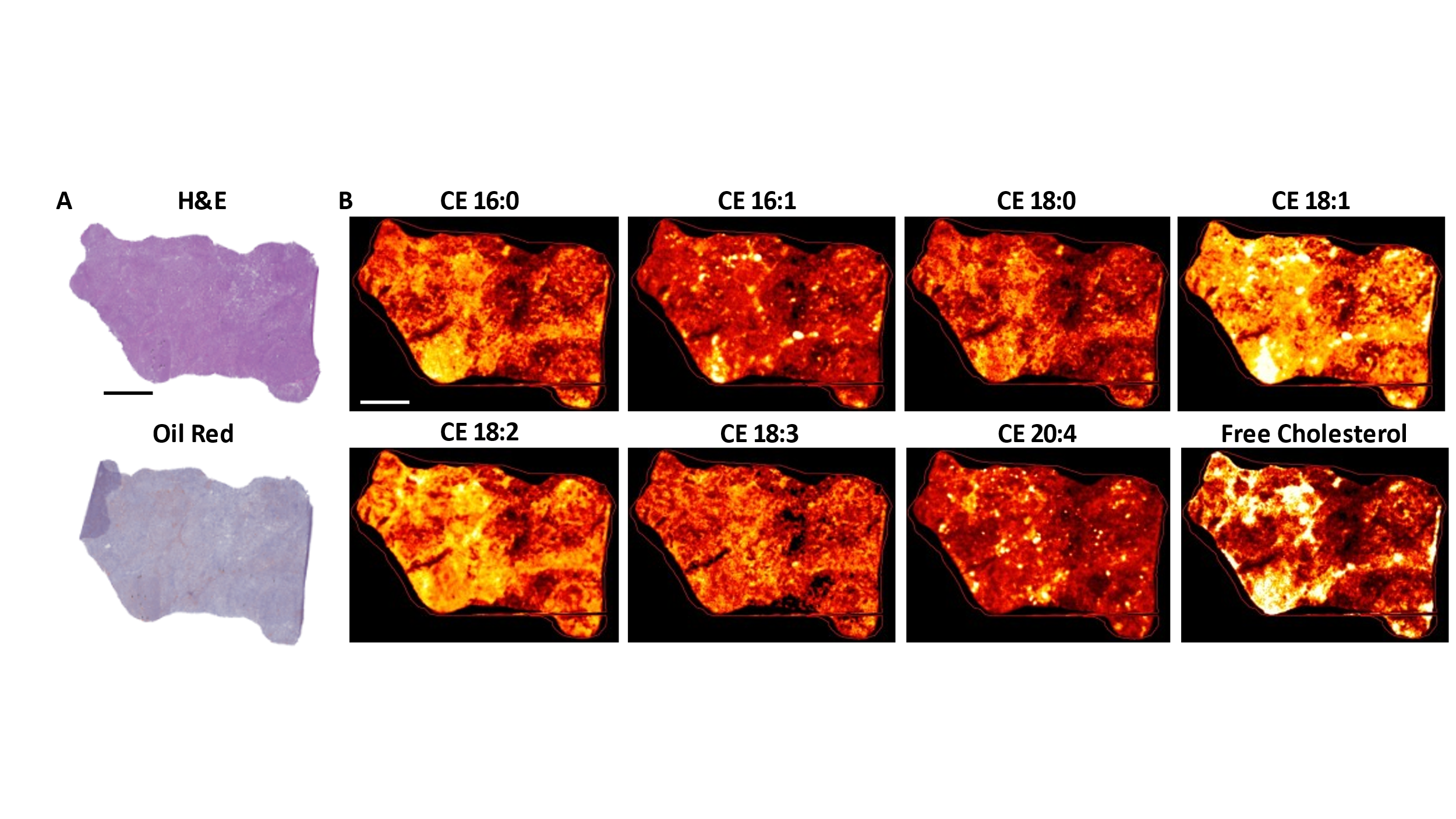

**Supplementary Fig. S3. Spatial distribution of cholesterol ester (CE) species in papillary renal cell carcinoma (pRCC) (patient P1). A.** Hematoxylin-eosin- and oil-red-stained kidney sections contiguous to those used for MALDI imaging. **B.** MALDI-2-MSI signal distribution for the major detected cholesterol ester species. Scale bar = 5 mm.

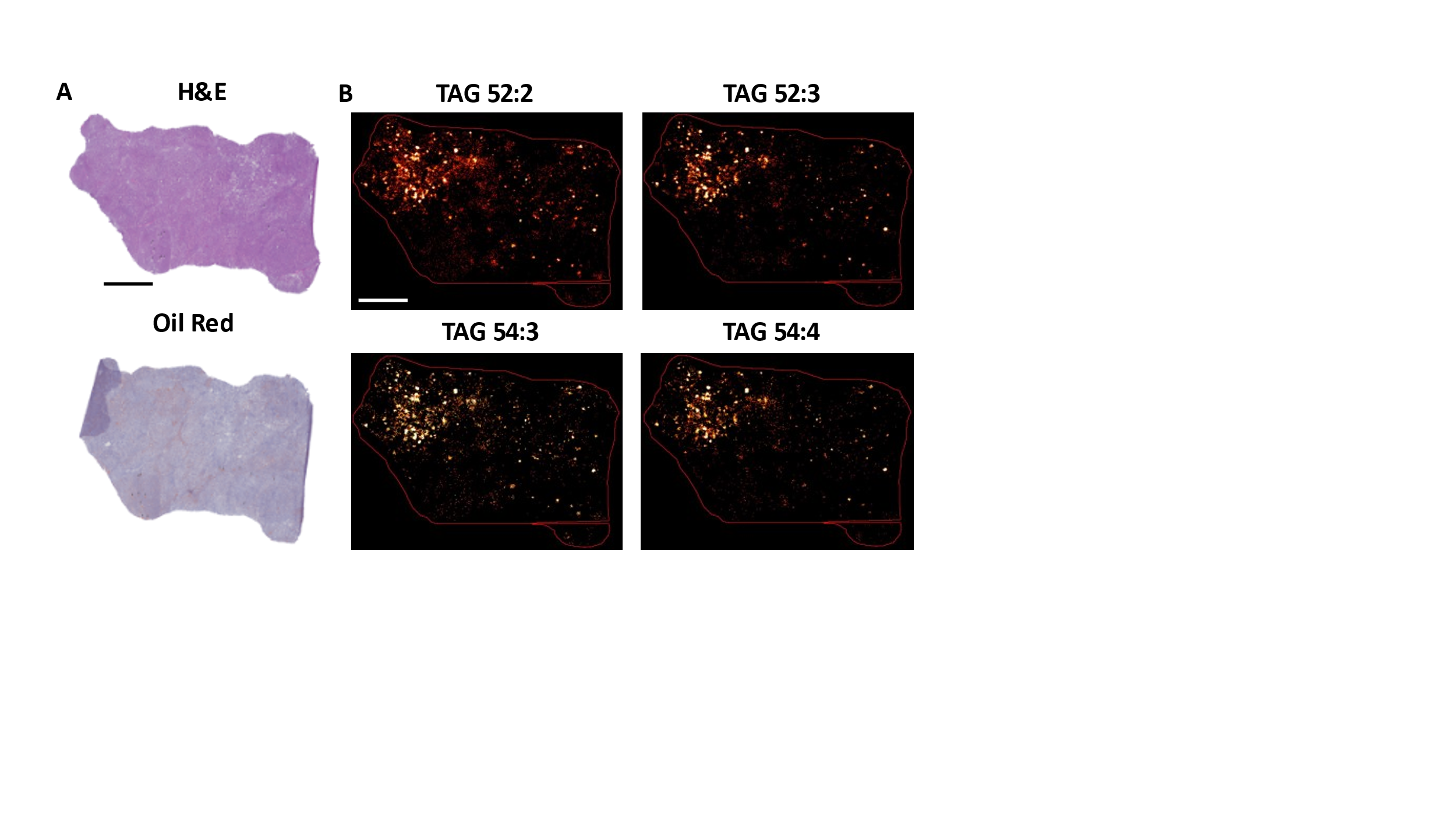

**Supplementary Fig. S4. Spatial distribution of triglyceride (TAG) species in papillary renal cell carcinoma (pRCC) (patient P1). A.** Hematoxylin-eosin- and oil-red-stained kidney sections contiguous to those used for MALDI imaging. **B.** MALDI-2-MSI signal distribution for four of the major detected TAG species. Scale bar = 5 mm

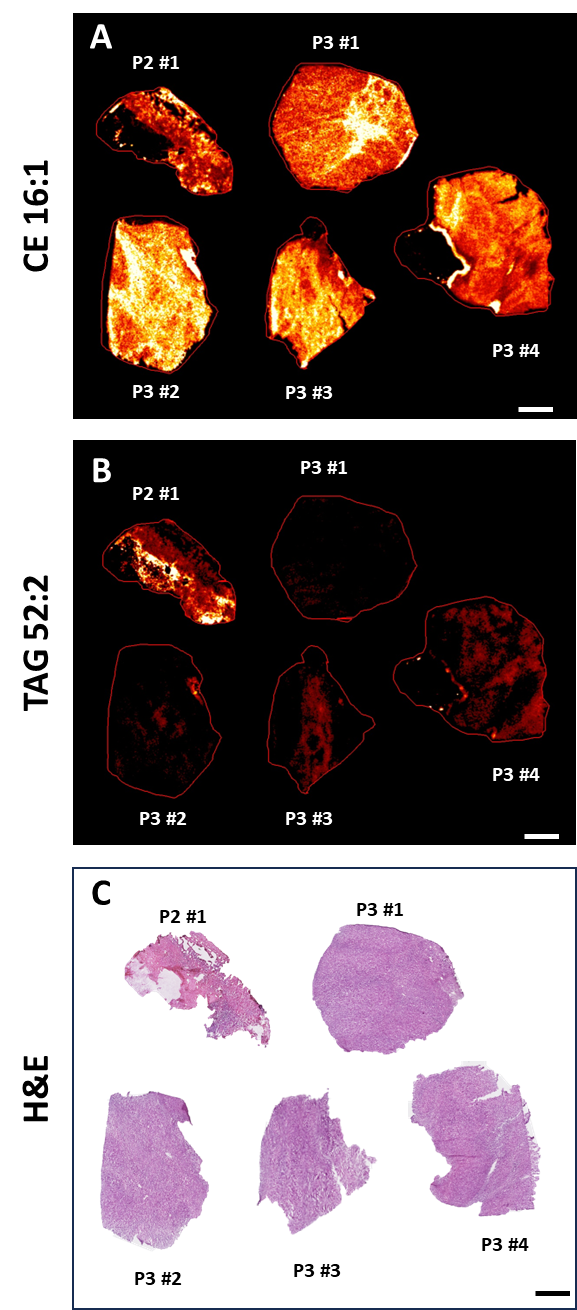

**Supplementary Fig. S5 Spatial distribution of triglyceride (TAG) and cholesteryl ester (CE) species in papillary renal cell carcinoma biopsies collected from patients P2 and P3. A.** MALDI-2-MSI signal distribution for CE 16:1. **B.** MALDI-2-MSI signal distribution for TAG 52:2. C. Hematoxylin-eosin-stained kidney sections contiguous to those used for MALDI imaging. P = patient ID, # = biopsy number. Scale bar = 5 mm

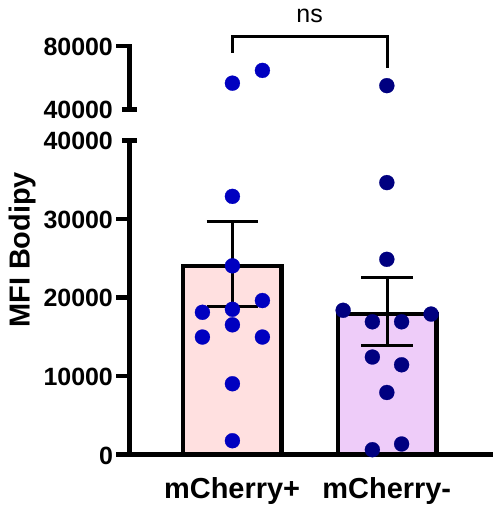

**Supplementary Fig. S6. Macrophage lipid droplet accumulation in *C. neoformans­-*infected cells.** Monocyte-derived macrophages (MDM) were infected with *C. neoformans* expressing mCherry and lipid droplet content was measured by imaging flow cytometry and expressed as median fluorescence intensity (MFI) of Bodipy 493/503, as described in Fig. 3. The figure shows Bodipy-associated fluorescence in macrophages carrying or not carrying intracellular fungi (mCherry+ and mCherry-) in the infected culture wells. Each dot represents one human donor. Mean and SEM are shown; ns, non-significant.

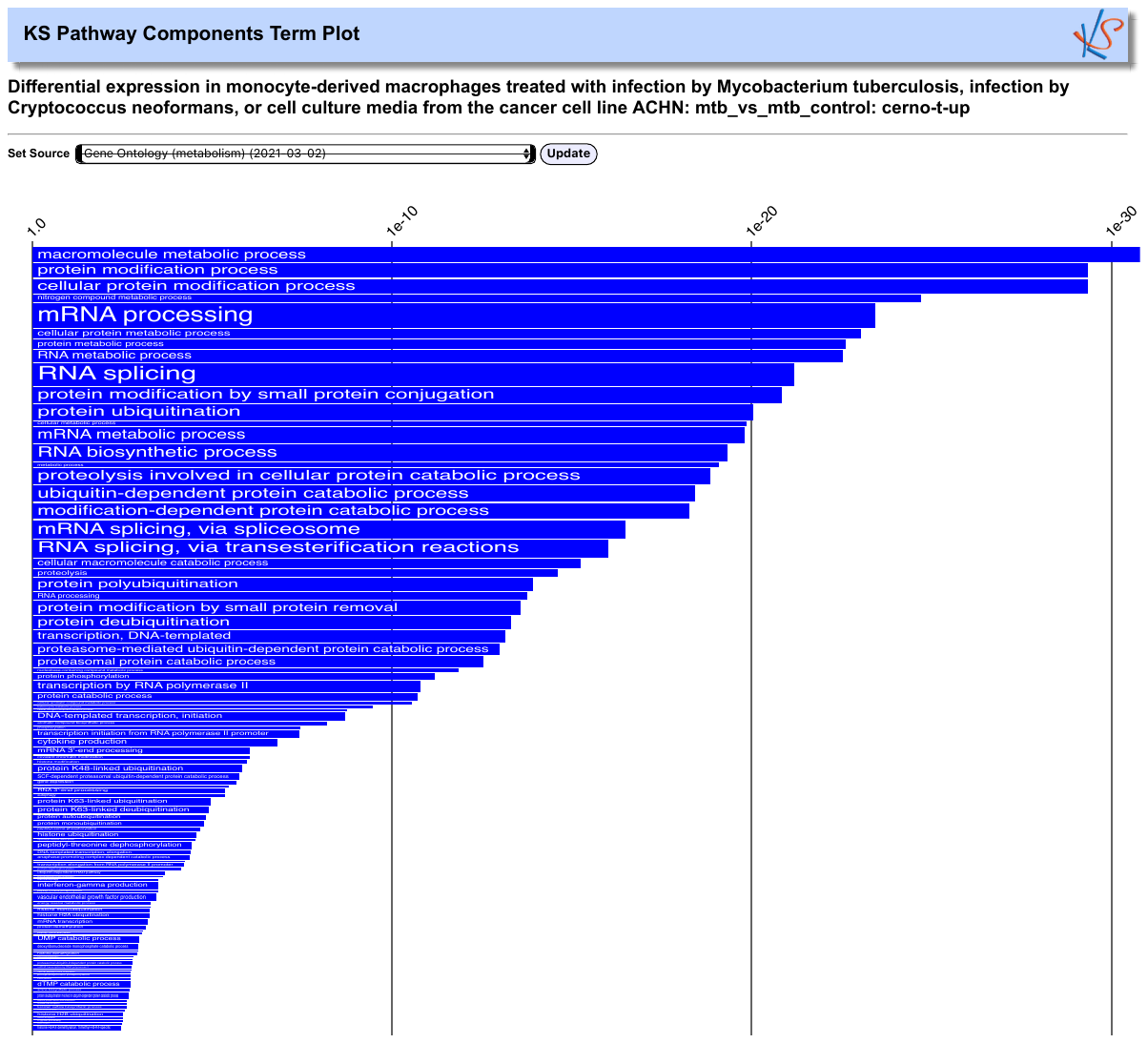

A. ***M. tuberculosis***

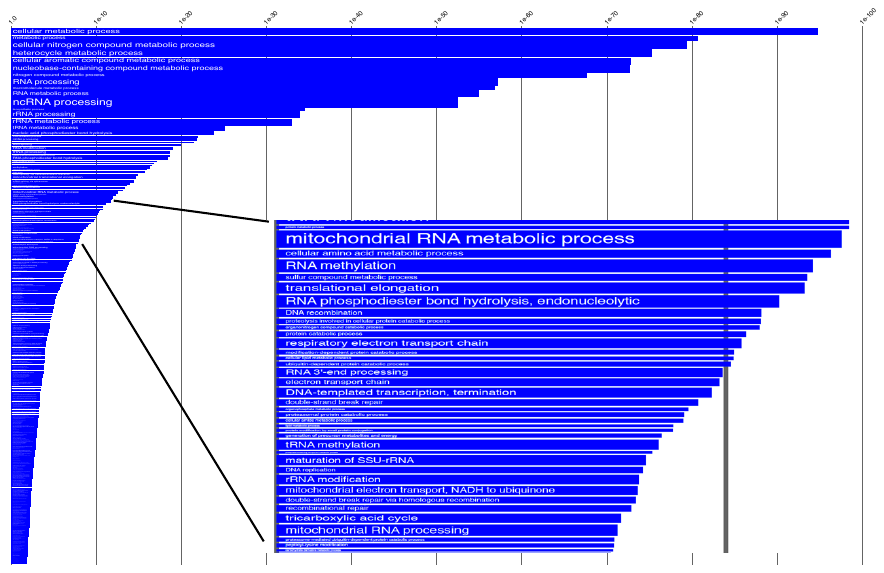

**B. *C. neoformans***

**Supplementary Fig. S7. A. Upregulated metabolism-associated pathways in *M. tuberculosis*-infected human monocyte-derived macrophages. B. Downregulated metabolism-associated pathways in *C. neoformans*-infected human monocyte-derived macrophages.** The panels show Gene Ontology (GO) annotations related to metabolic processes that were upregulated in *M. tuberculosis*-infected MDM and downregulated in *C. neoformans*-infected MDM, relative to uninfected control cells. The differential expression between sample classes (infected vs uninfected) was tested with coincident extreme ranks in numerical observations (CERNO). Pathways were selected using a cutoff false discovery rate of 0.05; the *p*-values for these pathways were plotted on the *x*-axis. To represent effect size, pathway gene sets containing fewer genes were given greater bar height/font size than were larger sets that yielded similar *p* values. For visualization purposes, panel A shows top-ranking annotations, and panel **B.** contains an inset highlighting mitochondria-related annotations.

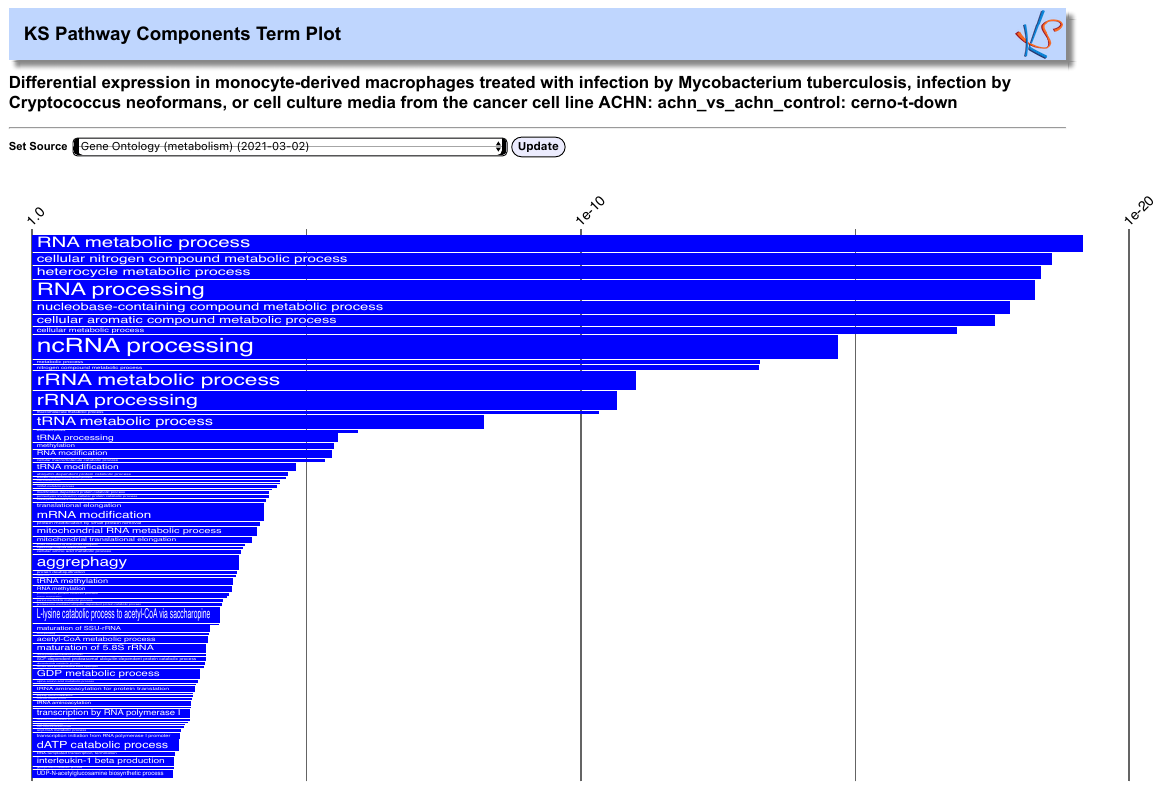
**Supplementary Fig. S8. Downregulated metabolism-associated pathways in ACHN-medium-treated human monocyte-derived macrophages.** The figure shows Gene Ontology (GO) annotations related to metabolic processes that were downregulated in MDM treated with ACHN conditioned medium, relative to untreated control cells. The differential expression between sample classes (treated vs untreated) was tested and selected pathways were represented as described in Figure S4. For visualization purposes, top-ranking annotations are shown.

**Supplementary Tables**

**Supplementary Table S1. List of triglyceride and cholesteryl ester species detected in tissues by MALDI-MSI.**

| **TAG** | **Detected in *Cryptococcus*** | **Detected in pRCC** | **CE** | **Detected in *Cryptococcus*** | **Detected in pRCC** |
| --- | --- | --- | --- | --- | --- |
| **Species** | **Tissue** | **Tissue** | **Species** | **Tissue** | **Tissue** |
| **48:0** | ✓ | 🗶 | **16:0** | ✓ | ✓ |
| **48:1** | ✓ | 🗶 | **16:1** | ✓ | ✓ |
| **48:2** | ✓ | 🗶 | **18:0** | ✓ | ✓ |
| **48:3** | ✓ | 🗶 | **18:1** | ✓ | ✓ |
| **50:0** | ✓ | 🗶 | **18:2** | ✓ | ✓ |
| **50:1** | ✓ | ✓ | **18:3** | ✓ | ✓ |
| **50:2** | ✓ | ✓ | **20:1** | ✓ | ✓ |
| **50:3** | ✓ | ✓ | **20:2** | 🗶 | ✓ |
| **50:4** | ✓ | 🗶 | **20:4** | ✓ | ✓ |
| **52:1** | ✓ | 🗶 |  |  |  |
| **52:2** | ✓ | ✓ |  |  |  |
| **52:3** | ✓ | ✓ |  |  |  |
| **52:4** | ✓ | ✓ |  |  |  |
| **52:5** | ✓ | ✓ |  |  |  |
| **52:6** | ✓ | 🗶 |  |  |  |
| **54:2** | ✓ | ✓ |  |  |  |
| **54:3** | ✓ | ✓ |  |  |  |
| **54:4** | ✓ | ✓ |  |  |  |
| **54:5** | ✓ | ✓ |  |  |  |
| **54:6** | ✓ | 🗶 |  |  |  |
| **54:7** | ✓ | 🗶 |  |  |  |
| **56:3** | ✓ | 🗶 |  |  |  |
| **56:4** | ✓ | 🗶 |  |  |  |

Triglyceride (TAG) species (identified by the total number of carbons and double bonds in the esterified fatty acids) and cholesteryl ester (CE) species detected by MALDI-MSI in the cryptococcus-infected lung tissues and papillary renal cell carcinoma (pRCC) tissue. ✓, detected; x, not detected.

**Supplementary Table S2a. Top five down- and up-regulated genes in macrophages infected with *M. tuberculosis.***

| **Gene ID** | **Gene** | **log2FoldChange** | ***p* value** |
| --- | --- | --- | --- |
| ENSG00000005469 | CROT | -0.694245982 | 6.81E-10 |
| ENSG00000183549 | ACSM5 | -1.875070163 | 2.15E-12 |
| ENSG00000072210 | ALDH3A2 | -0.675169083 | 1.18E-15 |
| ENSG00000131238 | PPT1 | -0.769679647 | 6.47E-16 |
| ENSG00000168765 | GSTM4 | -0.846851963 | 4.64E-14 |
| ENSG00000105325 | FZR1 | 0.451795 | 1.01E-10 |
| ENSG00000186187 | ZNRF1 | 0.884946 | 4.79E-17 |
| ENSG00000108306 | FBXL20 | 0.849817 | 9.11E-17 |
| ENSG00000186591 | UBE2H | 0.795385 | 3.10E-25 |
| ENSG00000187555 | USP7 | 0.206178 | 4.81E-06 |

Genes in differentially expressed metabolism-related Gene Ontology gene sets having FDR < 0.05 were selected based on individual *p* value (*p* << 0.05) and ranked by log2 fold change (infected vs non-infected) and number of occurrences in the differentially expressed gene sets. The top five down- and up-regulated genes were selected for further analysis.

**Supplementary Table S2b. Genes functionally related with the top five down- and up-regulated genes in macrophages infected with *M. tuberculosis*.**

| **Gene ID** | **Gene** | **log2FoldChange** | ***p* value** |
| --- | --- | --- | --- |
| ENSG00000215009 | ACSM4 | -1.65627 | 2.66E-05 |
| ENSG00000130377 | ACSBG2 | -1.09841 | 3.99E-03 |
| ENSG00000140284 | SLC27A2 | 0.929524 | 2.40E-03 |
| ENSG00000143554 | SLC27A3 | 0.440446 | 2.69E-02 |
| ENSG00000110090 | CPT1A | -0.69747 | 6.60E-05 |
| ENSG00000157184 | CPT2 | -0.32849 | 3.84E-05 |
| ENSG00000154227 | CERS3 | 1.207873 | 6.71E-03 |
| ENSG00000139624 | CERS5 | 0.437953 | 5.94E-06 |
| ENSG00000177076 | ACER2 | -0.46598 | 2.77E-02 |
| ENSG00000141510 | TP53 | -0.34037 | 1.19E-04 |
| ENSG00000109670 | FBXW7 | 0.514687 | 0.00000987 |
| ENSG00000135679 | MDM2 | 0.745793 | 1.91E-10 |
| ENSG00000100219 | XBP1 | 1.329735 | 1.36E-11 |
| ENSG00000155792 | DEPTOR | -2.04205 | 0.0000007 |
| ENSG00000164181 | ELOVL7 | 4.018986 | 1.04E-33 |
| ENSG00000175445 | LPL | -1.17544 | 6.57E-08 |
| ENSG00000134184 | GSTM1 | -0.95459 | 0.000000069 |
| ENSG00000213366 | GSTM2 | -1.23318 | 0.000000626 |
| ENSG00000166851 | PLK1 | -1.07326 | 0.0025 |
| ENSG00000092964 | DPYSL2 | -0.9294 | 1.38E-06 |
| ENSG00000013588 | GPRC5A | 6.429513 | 2.92E-81 |
| ENSG00000181773 | GPR3 | 3.096538 | 1.07E-34 |
| ENSG00000183484 | GPR132 | 3.238435 | 2.91E-26 |
| ENSG00000180758 | GPR157 | 2.357273 | 5.3E-21 |
| ENSG00000173698 | GPR64 | 2.313555 | 8.38E-17 |
| ENSG00000139572 | GPR84 | 2.131323 | 4.78E-10 |
| ENSG00000182885 | GPR97 | 2.623284 | 1.95E-09 |
| ENSG00000188316 | ENO4 | -1.32246 | 7.67E-07 |
| ENSG00000077721 | UBE2A | 0.552901 | 1.40E-08 |
| ENSG00000160087 | UBE2J2 | 0.405355 | 2.54E-08 |
| ENSG00000177889 | UBE2N | 0.348958 | 3.68E-08 |
| ENSG00000131508 | UBE2D2 | 0.331604 | 7.16E-07 |
| ENSG00000185651 | UBE2L3 | 0.381012 | 3.78E-06 |
| ENSG00000160714 | UBE2Q1 | 0.35768 | 2.01E-05 |
| ENSG00000072401 | UBE2D1 | 0.867935 | 3.27E-05 |
| ENSG00000104343 | UBE2W | 0.503851 | 4.69E-05 |
| ENSG00000182247 | UBE2E2 | 0.746942 | 0.000065 |
| ENSG00000078140 | UBE2K | 0.277765 | 0.000104 |
| ENSG00000170142 | UBE2E1 | 0.523719 | 0.000327 |
| ENSG00000156587 | UBE2L6 | 1.059927 | 0.000437 |
| ENSG00000175931 | UBE2O | 0.349526 | 0.000465 |
| ENSG00000130725 | UBE2M | 0.31568 | 0.000899 |
| ENSG00000107341 | UBE2R2 | 0.236068 | 0.002875 |
| ENSG00000184182 | UBE2F | 0.245525 | 0.021078 |
| ENSG00000198833 | UBE2J1 | 0.305753 | 0.02493 |
| ENSG00000187531 | SIRT7 | 0.5201 | 1.33E-10 |
| ENSG00000177463 | NR2C2 | 0.771918 | 0.000254 |
| ENSG00000182054 | IDH2 | -0.83851 | 2.92E-14 |
| ENSG00000138413 | IDH1 | -0.94364 | 2.62E-18 |
| ENSG00000142208 | AKT1 | -0.16012 | 0.045604 |
| ENSG00000082701 | GSK3B | 0.515329 | 0.000158 |

Differentially expressed genes (*p* < 0.05) reported in the published literature as functionally related to the top five up- and down-regulated genes in Table S2a.

**Supplementary Table S3a. Top five down- and up-regulated genes in macrophages infected with *C. neoformans.***

| **Gene** | **Gene Id** | **log2FoldChange** | ***p* value** |
| --- | --- | --- | --- |
| ENSG00000116750 | UCHL5 | -0.20727 | 0.04472 |
| ENSG00000138035 | PNPT1 | -0.54945 | 1.67E-10 |
| ENSG00000130348 | QRSL1 | -0.37168 | 3.88E-07 |
| ENSG00000196510 | ANAPC7 | -0.39341 | 0.000302 |
| ENSG00000131467 | PSME3 | -0.30045 | 3.30E-05 |
| ENSG00000109107 | ALDOC | 1.825971589 | 2.32E-10 |
| ENSG00000111674 | ENO2 | 1.489680497 | 9.70E-15 |
| ENSG00000159399 | HK2 | 1.121945578 | 6.74E-16 |
| ENSG00000102144 | PGK1 | 0.746990585 | 2.14E-08 |
| ENSG00000111640 | GAPDH | 0.713009241 | 1.06E-07 |

Gene identification for gene-level analysis was performed as in Table S2a.

**Supplementary Table S3b. Genes functionally related with the top five down- and up-regulated genes in macrophages infected with *C. neoformans.***

| **Gene** | **Gene Id** | **log2FoldChange** | ***p* value** |
| --- | --- | --- | --- |
| ENSG00000185920 | PTCH1 | 0.761480524 | 0.012629752 |
| ENSG00000127528 | KLF2 | 0.62022683 | 0.047195731 |
| ENSG00000166819 | PLIN1 | 0.973316711 | 0.003288899 |
| ENSG00000147872 | PLIN2 | 0.692199046 | 0.00059194 |
| ENSG00000149925 | ALDOA | 0.568357915 | 4.63E-06 |
| ENSG00000111669 | TPI1 | 0.6108752 | 5.94E-07 |
| ENSG00000171314 | PGAM1 | 0.429844596 | 0.001449782 |
| ENSG00000105220 | GPI | 0.684479515 | 7.26E-09 |
| ENSG00000067057 | PFKP | 0.690877949 | 0.000281329 |
| ENSG00000152256 | PDK1 | 1.070412291 | 8.18E-08 |
| ENSG00000067992 | PDK3 | 0.363881857 | 0.004868894 |
| ENSG00000134333 | LDHA | 1.050510273 | 3.90E-08 |
| ENSG00000104812 | GYS1 | 0.794960339 | 3.17E-07 |
| ENSG00000174437 | ATP2A2 | -0.330898147 | 0.029025124 |
| ENSG00000132356 | PRKAA1 | -0.186245418 | 0.045627492 |
| ENSG00000131828 | PDHA1 | -0.222577362 | 0.025045384 |
| ENSG00000072210 | ALDH3A2 | -0.331695382 | 7.63E-05 |
| ENSG00000161618 | ALDH16A1 | -0.281990927 | 0.008872148 |
| ENSG00000184254 | ALDH1A3 | -0.998071894 | 0.011142259 |
| ENSG00000159423 | ALDH4A1 | -0.573208906 | 0.013428796 |
| ENSG00000137124 | ALDH1B1 | -0.500558706 | 0.026687482 |
| ENSG00000135218 | CD36 | -0.435604275 | 0.045176474 |

Differentially expressed genes (*p* < 0.05) reported in the published literature as functionally related to the top five up- and down-regulated genes in Table S3a.

**Supplementary Table S4a. Top five down and upregulated genes in macrophages treated with ACHN-conditioned culture medium.**

| **Gene** | **Gene Id** | **log2FoldChange** | ***p* value** |
| --- | --- | --- | --- |
| ENSG00000109819 | PPARGC1A | -1.108179567 | 0.0019129 |
| ENSG00000179598 | PLD6 | -0.340258537 | 0.0313289 |
| ENSG00000164742 | ADCY1 | -1.789945859 | 1.59E-05 |
| ENSG00000156239 | N6AMT1 | -0.407562926 | 0.0036691 |
| ENSG00000215009 | ACSM4 | -0.732880759 | 0.0301056 |
| ENSG00000111674 | ENO2 | 0.585748 | 0.005669 |
| ENSG00000160883 | HK3 | 0.417557 | 0.035561 |
| ENSG00000111669 | TPI1 | 0.280168 | 0.035629 |
| ENSG00000102144 | PGK1 | 0.263218 | 0.041578 |
| ENSG00000162433 | AK4 | 0.550335 | 0.00025 |

Gene identification for gene-level analysis was performed as in Table S2a.

**Supplementary Table S4b. Genes functionally related with the top five down and upregulated genes in macrophages treated with ACHN conditioned culture medium.**

| **Gene** | **Gene Id** | **log2FoldChange** | ***p* value** |
| --- | --- | --- | --- |
| ENSG00000135218 | CD36 | -0.71398 | 0.002177 |
| ENSG00000166411 | IDH3A | -0.36846 | 0.012774 |
| ENSG00000110090 | CPT1A | -0.3713 | 0.048744 |
| ENSG00000165092 | ALDH1A1 | -0.6716 | 0.005934 |
| ENSG00000214456 | PLIN5 | 0.763578 | 0.033995 |
| ENSG00000007866 | TEAD3 | 0.442895 | 0.043061 |
| ENSG00000142583 | SLC2A5 | 0.948899 | 0.001353 |
| ENSG00000114268 | PFKFB4 | 0.682204 | 3.73E-05 |

Differentially expressed genes (*p* < 0.05) reported in the published literature as functionally related to the top five up- and down-regulated genes in Table S5a.

**Supplementary text on transcriptomics analyses**

1. **Metabolism-related gene-level analysis of *M. tuberculosis*-infected monocyte-derived macrophages.**

Genes in differentially expressed metabolism-related Gene Ontology gene sets having FDR < 0.05 were selected based on individual *p* value (*p* << 0.05) and ranked by log2 fold change (infected vs non-infected) and number of occurrences in the differentially expressed gene sets. We analyzed the top five down- and up-regulated genes (**Table S2a**) and differentially expressed genes (*p* < 0.05) that were functionally related to them (**Table S2b**). In the text below, and in the subsequent sections, the names of the top five up- and down-regulated genes (**Table S2a**) are shown in bold. The names of the up- and down-regulated functionally related genes (**Table S2b**) are underlined.

Multiple mechanisms of decreased fatty acid oxidation were identified. These included downregulation of genes involved in (i) synthesis of acyl-CoA (acyl-CoA synthases ACSM4, **ACSM5**, ACSBG2), (ii) acyl-CoA conversion to acyl-carnitine [(carnitine transferase in peroxisome (**CROT**) and mitochondria (CPT1A, CPT2)], which is required for transport across the mitochondrial membrane (1-3), and (iii) oxidation to fatty acids of fatty aldehydes and fatty alcohols derived from lipid catabolism (fatty aldehyde dehydrogenase, **ALDH3A2**) (4-6).

Gene expression markers of increased production of triacylglycerols (TAG) were also identified. One set of gene expression changes were associated with increased production of ceramide, which induces triglyceride production (7-10). These included (i) downregulation of the aldehyde hydrogenase **ALDH3A2**, which converts hexadecenal to hexadecenoic acid. The resulting hexadecenal accumulation may lead to ceramide production; (ii) downregulation of ceramidase ACER2, and (iii) upregulation of ceramide synthases (CERS3 and CERS5). An additional marker of triglyceride accumulation was the downregulation of the gene encoding palmitoyl-protein thioesterase 1 (**PPT1**), which expresses lipolytic activity (11, 12). TAG accumulation in *M. tuberculosis*-infected macrophages may also result from altered cellular redox. For example, we observed downregulation of genes encoding glutathione S-Transferase Mu (GSTM1, GSTM2, **GSTM4**), a scavenger of reactive oxygen species (ROS), which can induce lipid droplet formation by inducing the ASK1/p38/JNK pathway (13-15).

As mentioned in the main text, *M. tuberculosis*-infected macrophages exhibited downregulated TP53 gene and upregulated TP53-specific E3 ligases (FBXW7, MDM2) that target this factor for proteasomal degradation (16-18). Upregulation of XBP1 can also lead to decreased TP53 levels by inducing MDM2 (19). Reduced TP53 activity well correlates with downregulation of TP53-dependent Deptor, which negatively regulates mTOR (20, 21). mTORC1 signaling is lipogenic in multiple ways, including increased de novo lipid synthesis, reduced lipolysis, and reduced fatty acid oxidation (22, 23).

We also observed upregulation of metabolism-related ubiquitination processes, which was prominent both at the pathway- and gene-level (**Fig. S4** and **Table S2a**). At the gene level, the top five upregulated genes featured genes encoding E3 ligases (**ZNRF1**) and associated proteins (**FZR1**, **FBXL20**, **UBE2H**) and the deubiquitinase **USP7,** which increases ZNRF1 activity as well as MDM2 activity (24-29). These changes presumably result in substantial macrophage proteome remodeling in response to *M. tuberculosis* infection. In particular, they may affect a variety of lipid metabolism regulators, including AKT1 and PLK1 (30, 31). **USP7** also stabilizes sirtuins (e.g., SIRT7, which is also upregulated at the gene level), resulting in increased expression of factors associated with TAG synthesis (24, 32).

2. **Metabolism-related gene-level analysis of *C. neoformans*-infected monocyte-derived macrophages.**

The top five up- and down-regulated genes (**Table S3a**) and functionally related, differentially expressed genes (**Table S3b**) were determined as described above for *M. tuberculosis* infection.

The top five upregulated genes encode glycolytic enzymes, strongly indicating a prominent role of a switch to glycolysis in *C. neoformans*-infected macrophages, which leads to increased triglyceride biosynthesis in multiple ways. Increased expression of hexokinase 2 (**HK2**) and other glycolytic enzymes (GPI, PFKP) results in increased production of fructose 1,6-bisphosphate, which can be converted to dihydroxyacetone phosphate (DHAP) and glyceraldehyde 3-phosphate (G3P), due to the increased production of aldolases (**ALDOC**, ALDOA). Also, the upregulation of the triosephosphate isomerase-encoding gene (TPI) might lead to increased production of DHAP, which is a precursor for triglyceride biosynthesis. The upregulation of additional glycolytic genes (**GAPDH**, **PGK1**, PGAM1, **ENO2**) might result in increased synthesis of phosphoenolpyruvate (PEP) (33). Increased PEP levels might inhibit sarco/endoplasmic reticulum Ca2+-ATPase, resulting in calcium imbalance and endoplasmic reticulum stress, which ultimately lead to lipogenesis (34-36). Moreover, the products of glycolytic genes may favor lipid accumulation in ways other than their direct role in glycolysis. For example, the **HK2** product participates in the metabolic switch from oxidative phosphorylation to glycolysis (37) and inhibits lipases (38). In addition, phosphoglycerate kinase 1 (**PGK1**) can phosphorylate pyruvate dehydrogenase kinases, which are also upregulated (PDK1, PDK3), and inhibit the pyruvate dehydrogenase (PDH) complex (39). Inhibition of the PDH complex, further compounded by the downregulation of one of its subunit-encoding genes (PDHA1), reduces conversion of pyruvate to acetyl CoA, leading to impaired tricarboxylic acid cycle and enhanced glycolysis (40). As a result, pyruvate may accumulate and be converted to lactate by lactate dehydrogenase (LDHA) (41), which we find upregulated. Lactate may inhibit lipolysis, thus resulting in lipid accumulation (42).

As mentioned in the main text, we also found indicators of reduced mitochondrial functions in *C. neoformans*-infected macrophages, including downregulation of polyribonucleotide nucleotidyl transferase 1 (**PNPT1**) and a glutaminyl-tRNA amidotransferase subunit 1 (**QRSL1**). The **PNPT1** product regulates the expression of the electron transport chain components at the mRNA and protein levels. Thus, downregulation of this gene might result in reduced function of the respiratory chain and decreased oxidative phosphorylation, leading to lipid accumulation (43, 44). The **QRSL1** product enables glutaminyl-tRNA synthase (glutamine-hydrolyzing) activity in the mitochondria. Missense mutations in the human QRSL1 locus have been associated with defects in mitochondrial respiratory chain complex and oxidative phosphorylation (45, 46). The resulting oxidative stress enhances lipogenesis (47, 48).

Lipid accumulation in *C. neoformans*-infected macrophages may also result from dysregulation of various other cellular activities. First, infected cells may exhibit increased fatty acid synthesis and decreased fatty acid oxidation due to the downregulation of (i) AMP-activated protein kinase (AMPK/PRKAA1), which inhibits de novo biosynthesis of fatty acids, increases fatty acid uptake, and stimulates fatty acid oxidation (49-51) and (ii) multiple genes encoding aldehyde dehydrogenases (ALDH3A2, ALDH16A1, ALDH1A3, ALDH4A1, ALDH1B1), which are involved in fatty acid oxidation (6). Second, perilipin genes (PLIN1, PLIN2), which are structural components of lipid droplets (52), are upregulated. PLIN gene upregulation may be associated, at least in part, with the downregulation of proteasome activator subunit 3 (**PSME3**) (53). The PSME3 product induces the degradation of Kruppel Like Factor 2 (KLF2) (54). Since KLF2 promotes the transcription of TAG synthesis and lipid storage genes, including PLIN1 and PLIN2 (55), increased KLF2 activity secondary to PSME3 downregulation is associated with lipid accumulation (56). Third, one may hypothesize that another system involved in *C. neoformans*-infection-induced lipid accumulation is Hedgehog signaling. Inhibition of Hedgehog signaling results in fat accumulation (57, 58). Downregulation of ubiquitin C-terminal hydrolase L5 (**UCHL5**), which positively regulates Hedgehog signaling by deubiquitinating one of its components, the Smoothened (SMO) protein (59, 60), might result in reduced Hedgehog signaling and increased lipid accumulation. The observed upregulation of patched1 (PTCH1), which represses SMO (61, 62), is consistent with a role of Hedgehog in lipid accumulation in *C. neoformans*-infected macrophages. Fourth, the downregulation of anaphase promoting complex subunit 7 (**ANAPC7**) may also lead to triglyceride accumulation. ANAPC7, which is a component of the anaphase promoting complex/cyclosome (APC/C) (63), interacts with the CBP/p300 coactivator and stimulates its acetyl transferase activity (64). Among the CBP/p300 targets is farnesoid X receptor (FXR) (65). Acetylation of FXR leads to protein stabilization but reduces its activity as transcriptional regulator (66). Thus, reduced FXR acetylation resulting from ANAPC7 downregulation might activate FXR signaling. FXR signaling may induce lipid accumulation and reduced fatty acid oxidation (67-69). However, some reports show the opposite effect of FXR signaling on lipid accumulation (70, 71), suggesting perhaps context-specific effects.

3. **Metabolism-related gene-level analysis of monocyte-derived macrophages treated with ACHN culture medium.**

The top five up- and down-regulated genes (**Table S4a**) and functionally related, differentially expressed genes (**Table S4b**) were determined as described above for *M. tuberculosis* infection.

The top upregulated genes are enriched for glycolytic genes. Increased expression of hexokinase 3 (**HK3**) and other glycolytic genes (PFKFB4) might result in increased production of fructose 1,6-bisphosphate, which breaks down into dihydroxyacetone phosphate (DHAP) and glyceraldehyde 3-phosphate (G3P) (37, 72). Upregulation of triosephosphate isomerase (**TPI1**) also contributes to the conversion of DHAP to G3P, favoring triglyceride synthesis (73). G3P formation can also lead to synthesis of phosphoenolpyruvate (PEP), due to upregulation of other glycolytic genes (**PGK1**, **ENO2**) (33). Accumulation of PEP may result in enhanced production of pyruvate. Reduced utilization of pyruvate in the TCA cycle, which may be downregulated (see below), might lead to enhanced lipogenesis (34).

Macrophages treated with ACHN medium also exhibit transcriptional markers of reduced production/activity of the energy sensor AMP-activated protein kinase (AMPK). One is the upregulation of adenylate kinase 4 (**AK4**), which regulates cellular ATP levels and inhibits AMPK signaling (74). Another is the downregulation of phospholipase D family member 6 (**PLD6**), which activates AMPK by altering mitochondrial fusion and fission dynamics (75). The resulting decreased activity of AMPK, an energy sensor maintaining energy homeostasis (76), might result in increased YAP/TAZ signaling, which regulates metastasis and metabolic reprogramming in cancer cells (77), since AMPK inhibits YAP/TAZ via YAP phosphorylation (78)**. This scenario is suggested by the observed upregulation of genes associated with or regulated by YAP/TAZ, such as TEAD3, SLC2A5, and PLIN5, in ACHN-medium-treated macrophages.** YAP/TAZ interacts with the TEAD transcription factors (79) to induce various target genes. One is the fructose transporter SLC2A5; its upregulation might result in increased fructose uptake, leading to lipogenesis via fructolysis (80, 81). YAP/TAZ also induces PLIN5, a member of a family of lipid-droplet-associated proteins (82), which protects triglycerides from enzymatic degradation by inhibiting adipose triglyceride lipase (83, 84). More in general, YAP/TAZ signaling and lipid metabolism are intricately intertwined in cancer cells, and YAP/TAZ is involved in the metabolism of fatty acids and sterols (77)**, suggesting a role of this signaling pathway also in the cholesterol accumulation detected in the ACHN-medium-treated cells.**

We identified additional gene expression markers of lipid accumulation. Impaired fatty acid oxidation, which may result in lipid accumulation, is suggested by the downregulation of the acyl-CoA synthase **ACSM4** and functionally related genes, such as acyl-CoA conversion to acyl-carnitine (carnitine palmitoyl transferase I, CPT1) and oxidation of fatty aldehydes to fatty acids (aldehyde hydrogenases, ALDH genes) (1-3, 6). The downregulation of **N6AMT1**, which encodes a methyltransferase mostly implicated in DNA and protein modification, is also consistent with lipid accumulation, since this gene product has been associated with lipid catabolism and low triglyceride levels (85, 86) by yet unknown mechanisms.

One of the top five downregulated genes is peroxisome proliferator-activated receptor gamma coactivator 1-alpha (**PPARGC1a**), a transcriptional coactivator that regulates cellular metabolism (87). Downregulation of PPARGC1a is consistent with the observed downregulation of two of its target genes, NAD-dependent isocitrate dehydrogenase 3 catalytic subunit alpha (IDH3A) (88) and CD36. Isocitrate dehydrogenases catalyze the oxidative decarboxylation of isocitrate to 2-oxoglutarate, which is a rate-limiting step of the TCA cycle. Thus, IDH3A downregulation might result in TCA cycle disruption. An additional indicator of dampened TCA cycle is the above-mentioned upregulation of **AK4**, since this adenylate kinase is associated with decreased expression of key TCA cycle genes (89). The scavenger receptor CD36 is a critical fatty acid sensor and regulator of lipid metabolism (90, 91). CD36 mediates fatty acid uptake, increases cholesterol efflux, and favors proteasomal degradation of HMG-CoA reductase, the rate-limiting enzyme in sterol synthesis (92, 93). CD36 downregulation might result in cholesterol accumulation, reduced fatty acid oxidation and mitochondrial function, and increased fatty acid storage (92). Thus, CD36 downregulation is consistent with increased accumulation of both cholesterol and triglycerides in ACHN-medium-treated cells. The decreased activity of AMPK proposed above would further decrease CD36 function, since AMPK promotes CD36 expression and translocation (50).

An additional marker of dysregulated cholesterol homeostasis is the downregulation of adenylate cyclase (**ADCY1**). In response to G-protein signaling, adenylate cyclases are activated and generate cyclic AMP (cAMP) (94). cAMP participates in various signal transduction mechanisms regulating adipogenesis, lipolysis, and cholesterol efflux (94, 95). Reduced cAMP production may be associated with decreased cholesterol efflux and decreased lipolysis (94-97).

**Materials and Methods**

**Cell cultures.** Peripheral blood mononuclear cells (PBMC) were isolated and monocyte-derived macrophages (MDM) were generated as previously described (98). Briefly, human blood was obtained from the New York Blood Center (Long Island City, NY, USA), and PBMC were isolated by Ficoll density gradient centrifugation (Ficoll-Paque, GE Healthcare, Uppsala, Sweden). Isolated PBMC were washed and resuspended in serum-free RPMI-1640 medium (Corning, Manassas, VA, USA) supplemented with 4 mM L-glutamine (Corning, Manassas, VA, USA) and seeded at a density of 1 × 10^7^ cells/ml in tissue culture multiwell plates or flasks. After 4 hr, non-adherent cells were removed by washing five times with 1× PBS (Corning, Manassas, VA, USA) and the adherent fraction was cultured over 7 days in complete RPMI medium (RPMI-1640 supplemented with 10% fetal bovine serum (Seradigm, Radnor, PA, USA) and 4 mM L-glutamine.

The ACHN cell line was obtained from the American Type Culture Collection (ATCC). To obtain conditioned medium, cells were grown in complete RPMI for 6-9 passages in humidified atmospheric air containing 5% CO_2_ at 37°C. Cell culture medium was collected 24 hr after the last passage, filtered through a 0.4 μM filter to remove any cell debris, and stored in aliquots at -80°C.

**Pathogens.** mCherry-expressing *M*. *tuberculosis* H_37_Rv strain was grown in liquid medium to mid-log phase and frozen stocks prepared for macrophage infection as described (5). mCherry-expressing *C. neoformans* H99 strain was grown in YPD medium overnight at 30°C with constant agitation, washed three times with 1× PBS, and resuspended in 1× PBS before use in macrophage infection. The final fungal concentration was adjusted to 1 x 10^8^ cell/ml with complete RPMI medium [RPMI-1640 supplemented with 10% heat-inactivated fetal bovine serum (Seradigm, Radnor, PA, USA) and 4 mM L-glutamine]. A suspension of heat-killed *C. neoformans* was prepared by resuspending the fungal cells in 1× PBS and incubating the suspension at 75°C for 1 hr in a heat block. To prepare cell-free culture supernatant, *C. neoformans* H99 was grown in complete RPMI medium for 24 hr (end of logarithmic growth phase), filtered through a 0.2 μM filter, and used for macrophage treatments.

**Macrophage treatments**. For infection experiments, following 7 days of MDM differentiation, MDM were counted and infected with either *M*. *tuberculosis or C. neoformans*. The pathogen inoculum for infection was prepared by adding the microorganisms to supplemented RPMI-1640 medium (as above) to obtain an MOI of 4 colony-forming units (cfu) per cell. Pathogen clumps were disrupted by vortexing with sterile 3-mm-diameter glass beads for 2 min; this suspension was used for MDM infection. MDM were incubated with *M*. *tuberculosis* for 24h. MDM were infected with *C. neoformans* for 3 h, washed with 1× PBS three times, and incubated with fresh medium for 24 hr. When chemical inhibitors were tested, they were added to the MDM at the time of infection. For treatment of uninfected MDMs with recombinant TNFα, the cytokine was incubated with the MDMs for 24 hr.

At 24 hr post-infection, supernatants were collected and frozen for cytokine analysis and MDM were washed with 1× PBS three times and collected for subsequent analysis.

For treatment with ACHN conditioned medium or IL-8 recombinant cytokine (Abcam, Cambridge, England), after PBMC isolation, the adherent fraction of PBMC was treated with conditioned medium (50% fresh medium and 50% conditioned medium) or IL-8 for 7 days. Medium was replaced at 3 and 6 days of incubation. At day 7 post-treatment, supernatants were collected and frozen for cytokine analysis, and MDM were washed with 1× PBS three times and collected for subsequent analysis.

Chemical inhibitor doses were selected based on available EC_50_ data and toxicity profiles with untreated macrophages (Trypan Blue staining or MTS assay; CellTiter 96 Aqueous One Solution Cell Proliferation Assay Promega, Madison, WI, USA). Only inhibitor doses resulting in >90% cell viability were utilized. The following concentrations were used: 30-90 nM DGAT inhibitor A922500 (PubChem CID: 24768261) (Santa Cruz Biotechnology, Dallas, TX, USA), 0.4 nM rapamycin (mTORC1 inhibitor) (Selleckchem, Houston, TX, USA), and 10 µM ACAT inhibitor CAS 615264-52-3 (PubChem CID: [10019206](https://pubchem.ncbi.nlm.nih.gov/compound/10019206)) (Santa Cruz Biotechnology, Dallas, TX, USA).

**Imaging flow cytometry.** Imaging flow cytometry of MDM was performed as previously described (98). MDM were detached from tissue culture plates by incubating with 5 mM EDTA in 1× PBS pH 8 for 30 min followed by gentle scraping, washed once with 1× PBS, and fixed with 4% paraformaldehyde in 1× PBS for 45 min at room temperature. Cells were then washed with 1× PBS containing 0.1% bovine serum albumin (PBS-BSA), resuspended in 50 μl of PBS-BSA containing 5 μl of Fc receptor blocking solution, FcX (BioLegend, San Diego, CA), and incubated at room temperature for 5 min. After incubation, 50 μl of PBS-BSA containing 2.5 μl of CD11c APC antibody (clone S-HCL-3) or 5 μl of CD11c BV421 antibody (clone B-ly6 RUO) (BD Biosciences) were added to each tube, and samples were incubated at 4°C for 30 min. After washing with PBS-BSA, cells were stained with 0.3 μg/ml Bodipy 493/503 (Life Technologies, Carlsbad, CA) in 1× PBS for 15 min. For each experimental condition, data from 5,000–10,000 CD11c+ cells were acquired with an ImageStream^X^Mark II imaging flow cytometer (Amnis Corporation, Seattle, WA) using 60× magnification. Image data were analyzed by IDEAS software version 6.0 (Amnis Corporation, Seattle, WA) after applying a compensation matrix and selecting the region of interest (lipid droplets) with the Spot Mask tool. Median fluorescence intensity and spots per cell were extracted.

**Lipid quantification in macrophages.** MDM were detached from the cell culture plates as described above and transferred to microcentrifuge tubes. Cell pellets were obtained by culture centrifugation at 300*x g* and frozen for subsequent lipid measurement. Triglyceride-Glo™ Assay and Cholesterol/Cholesterol Ester-Glo™ Assay kits (Promega; Madison, WI, USA) were used to quantify triglycerides and cholesterol, following the manufacturer’s instructions. Cholesterol was quantified following addition of cholesteryl esterase to convert cholesteryl esters to free cholesterol, following manufacturer’s instructions.

**Mouse infection and tissue collection.** Female mice with an average weight of 20-25 g were used. C57BL/6 mice were purchased from the Jackson Laboratories. Animal studies were performed at the Public Health Research Institute Animal facility. All studies were conducted following biosafety level 2 (BSL-2) protocols and procedures approved by the Institutional Animal Care and Use Committee (IACUC) and Institutional Biosafety Committee of Rutgers University under protocol 999901066. *C. neoformans* wild type strain H99 was cultured on YPD medium. To prepare fungal cells for infection, an overnight culture of *C. neoformans* H99 was washed three times with 1× PBS and the concentration of yeast cells was determined by hemocytometer counting. The final fungal concentration was adjusted with PBS to 2 x 10^6^ cell/ml. Each mouse was infected intranasally with 1 x 10^5^ H99 cells in a 50 μl volume after being anesthetized with a mix of Ketamine (12.5 mg/mL) and Xylazine (1 mg/mL). After infection, animals were weighed daily and monitored twice daily for progression of disease, including weight loss, gait changes, labored breathing, and fur ruffling. Infected animals were sacrificed at day 7 post-infection, according to the experimental design and the Rutgers University IACUC approved animal protocol. A part of the dissected lung tissues was fixed in 10% formalin solution and sent to the Rutgers histopathological core facility for section preparation and staining with hematoxylin and eosin (H&E) using standard protocols. Another portion of the lung tissues was placed on labeled plastic cryohistology trays and frozen on dry ice for 2 minutes in a Styrofoam container. The samples were then wrapped with aluminum foil and stored in labeled Ziploc bags at -80°C until use for MALDI imaging.

**Human Biopsies.** Papillary renal cell carcinoma tissue was obtained from the Rutgers CINJ Biorepository. Tissue collected during surgery was frozen by the Biorepository using the PrestoCHILL machine, which allows for ultrafast freezing. All tissues analyzed derived from radical nephrectomies; the RWJUH surgical pathology reports indicated papillary renal cell carcinoma, type 1, WHO grade 2. Utilization of archived de-identified biospecimens from the Rutgers Cancer Institute of New Jersey Biospecimen Repository Services (BRS) Shared Resources was performed under IRB protocols 001006 and 002002.

**Matrix-assisted laser desorption/ionization (MALDI) mass spectrometry**. Frozen lung samples were used to prepare 12 μm-thick cryosections using a Leica CM1860 cryostat. Tissue sections were mounted onto indium tin oxide coated glass slides (Delta Technologies Limited, Loveland, Colorado) and stored individually in Ziploc bags at -80°C. Adjacent sections were prepared on Superfrost Plus™ glass microscope slides (Fisherbrand) for H&E staining, which was performed using standard protocols at the UTMB Histology core. Sections for MALDI imaging were coated with 2′,4′,6′-Trihydroxyacetophenone monohydrate at 10 mg/ml in 50:50 cyclohexane/methanol using a HTX TM Sprayer. 20 passes were performed over each tissue at a spray volume of 50 ml/min and nozzle temperature of 50°C. Once sprayed, samples were placed individually in Ziploc bags and immediately transferred to a -80°C freezer for storage. Following MALDI-MS image acquisition, matrix was removed from the slides by serial rinses in 50% ethanol, and tissue sections were allowed to airdry at room temperature for 30 min. The tissue sections were then transferred to the UTMB histology core for H&E and Oil Red staining using standard protocols.

MALDI MS imaging was performed using a Q-Exactive HF mass spectrometer (Thermo Scientific, Bremen, DE) fitted with a MALDI/ESI Injector (Spectroglyph LLC, Kennewick, WA). Laser post-ionization (MALDI-2) was used to enhance analytical sensitivity for triglycerides and cholesteryl esters. Images were acquired at 20 micrometer voxel size, using a pulse energy of ~6 mJ and repetition rate of 30Hz. Q Exactive HF MS Scan parameters were optimized for triglyceride and cholesteryl ester detection: polarity – positive, scan range – 350-1500 m/z, resolution – 120,000, automatic gain control - off, maximum inject time 250 ms**.** ImageInsight™ (Spectroglyph LLC) software was used for initial data visualization and to convert data files into imzML format for visualization and further processing in SCiLS™ software (Bruker, Billerica, MA). TAG and CE species were predominantly detected as potassium adducts and lipid identifications were assigned within a 3pm mass tolerance range using Lipid Maps database. All lipid images produced were normalized to the total ion chromatogram.

**Transcriptomics and pathway analysis.** RNA was isolated by using TRI reagent (Molecular Research Center, Cincinnati, OH, USA) according to manufacturer’s instructions. RNA-Seq was conducted at the Rutgers Genomics Center. The quality of RNA was first assessed for integrity on an Agilent TapeStation using high sensitivity RNA kit, PN-5067-5579 (Agilent technologies Inc, CA). Samples with RNA integrity number (RIN) >7.0 were considered to have sufficient quality for subsequent processing. Illumina compatible cDNA libraries generated for polyA selected mRNA using NEB next ultra RNA library preparation kit Cat#E7530L (New England Biolabs Inc, MA). The cDNA libraries were purified using AmpureXP beads, Product No: A63882 (Beckman Coulter) and analyzed on an Agilent TapeStation using HS D1000 screen tape PN- 5067-5584 (Agilent Technologies Inc, CA) to estimate the size of the library and quantitated using Qubit 4 Fluorometer, using HS reagent kit, Cat. No. Q33231 (Thermofisher Scientific, MA). Equimolar amounts of barcoded libraries pooled together and sequenced on Illumina NovaSeq 6000 Instrument (Illumina, San Diego, CA) using SP flow cells kits (cat # 20040326) with 2x150 cycles configuration.

Read count and gene-level transcript abundance analysis were performed by the Rutgers Molecular and Genomics Informatics Core. In brief, raw transcriptome reads were assessed for quality control (FASTQC v0.11.8) and trimmed for quality and adapter contaminant (cutadapt v 2.5). Trimmed reads were aligned to the mouse genome (GRCh37) using STAR (v2.6.1), followed by transcript abundance calculation and hit count extraction with StringTie (v2.0) and featureCounts (v1.6.4), respectively.

The differential expression between sample classes (treated vs non treated for each condition) was tested with coincident extreme ranks in numerical observations (CERNO) (99) for metabolism-related Gene Ontology gene sets. The Benjamini–Hochberg method was used to calculate the false discovery rate (FDR). Gene sets (pathways) were identified at a cutoff false discovery rate of 0.05. Genes in differentially expressed metabolism-related Gene Ontology gene sets having FDR < 0.05 were selected based on individual *p* value (*p* << 0.05) and ranked by log2 fold change (treated vs non-treated in each condition) and number of occurrences in the differentially expressed gene sets. The top five down- and up-regulated genes were selected for further analysis. Differentially expressed genes (*p* < 0.05) reported in the published literature as functionally related to the top five up- and down-regulated genes in each condition were also selected for analysis and data interpretation.
